# Scientific publications and COVID-19 “research pivots” during the pandemic: An initial bibliometric analysis

**DOI:** 10.1101/2020.12.06.413682

**Authors:** Philip Shapira

## Abstract

An examination is presented of scientific research publication trends during the global coronavirus (COVID-19) pandemic in 2020. After reviewing the timing of the emergence of the pandemic in 2020 and the growth of governmental responses, available secondary sources are used to highlight impacts of COVID-19 on scientific research. A bibliometric analysis is then undertaken to analyze developments in COVID-19 related scientific publications through to October of 2020 by broad trends, fields, countries, and organizations. Two publication data sources are used: PubMed and the Web of Science.

While there has been a massive absolute increase in PubMed and Web of Science papers directly focused on COVID-19 topics, especially in medical, biological science, and public health fields, this is still a relatively small proportion of publication outputs across all fields of science. Using Web of Science publication data, the paper examines the extent to which researchers across all fields of science have pivoted their research outputs to focus on topics related to COVID-19. A COVID-19 research pivot is defined as the extent to which the proportion of output in a particular research field has shifted to a focus on COVID-19 topics in 2020 (to date) compared with 2019. Significant variations are found by specific fields (identified by Web of Science Subject Categories). In a top quintile of fields, not only in medical specialties, biomedical sciences, and public health but also in subjects in social sciences and arts and humanities, there are relatively high to medium research pivots. In lower quintiles, including other subjects in science, social science, and arts and humanities, low to zero COVID-19 research pivoting is identified.

*In a new Appendix to the paper, an updated analysis is provided through to mid-April 2022*.

**Citation:** Shapira, P. “Scientific publications and COVID-19 “research pivots” during the pandemic: An initial bibliometric analysis,” *bioRxiv* 2020.12.06.413682; doi: https://doi.org/10.1101/2020.12.06.413682

**Version Notes:** Version 1: Original paper, completed on December 6, 2020; posted at *bioRxiv* on December 7, 2020.

Version 2: Minor grammar items corrected.

Version 3: Updated bibliometric analysis through to mid-April 2022 added on April 29, 2022, as new Appendix 2.

## Introduction

This working paper takes an initial look at scientific research publication trends during the global coronavirus (COVID-19) pandemic.

To provide context, the paper first discusses the timing of the emergence of the pandemic in 2020 and the growth of governmental responses. Scientific research, as with other societal and economic activities across the globe, has been greatly impacted by COVID-19.

As one approach (of many possible approaches) to assessing COVID-19’s effects on science, the paper analyzes developments in COVID-19 and all scientific publications in 2020 by broad trends, fields, countries, and organizations. Comparisons are made with the pre-COVID year of 2019. The paper also examines the extent to which researchers across all fields of science have pivoted their research outputs to focus on topics related to COVID-19.

Two publication data sources are used for the analyses presented in the paper: PUBMED and the Web of Science. A bibliometric search approach is applied to publication datasets from these two sources to identify COVID-19 relevant publications. The publication datasets and bibliometric search approach are described in a subsequent section of the paper.

### Emergence of the pandemic and timing of government responses

In January 2020, the COVID-19 outbreak was declared a public health emergency of international concern by the World Health Organization (WHO).^1^ In March 2020, WHO designated COVID-19 as a pandemic.^2^ While early cases of COVID-19 were reported in January 2020, the worldwide rate of growth of reported cases began to accelerate from February 2020 (Figure 1). Reported COVID-19 cases continued to increase globally throughout 2020, reaching 45.7 million by the end of October 2020, and growing to over 61 million by the end of November 2020.^3^ The number of countries reporting COVID-19 cases rose to 22 by the end of January 2020, rising to 58 by the end of February 2020. Just a month later, by the end of March 2020, COVID-19 cases were reported in 192 countries, rising to 212 by October 2020.^4^

**Figure 1.**
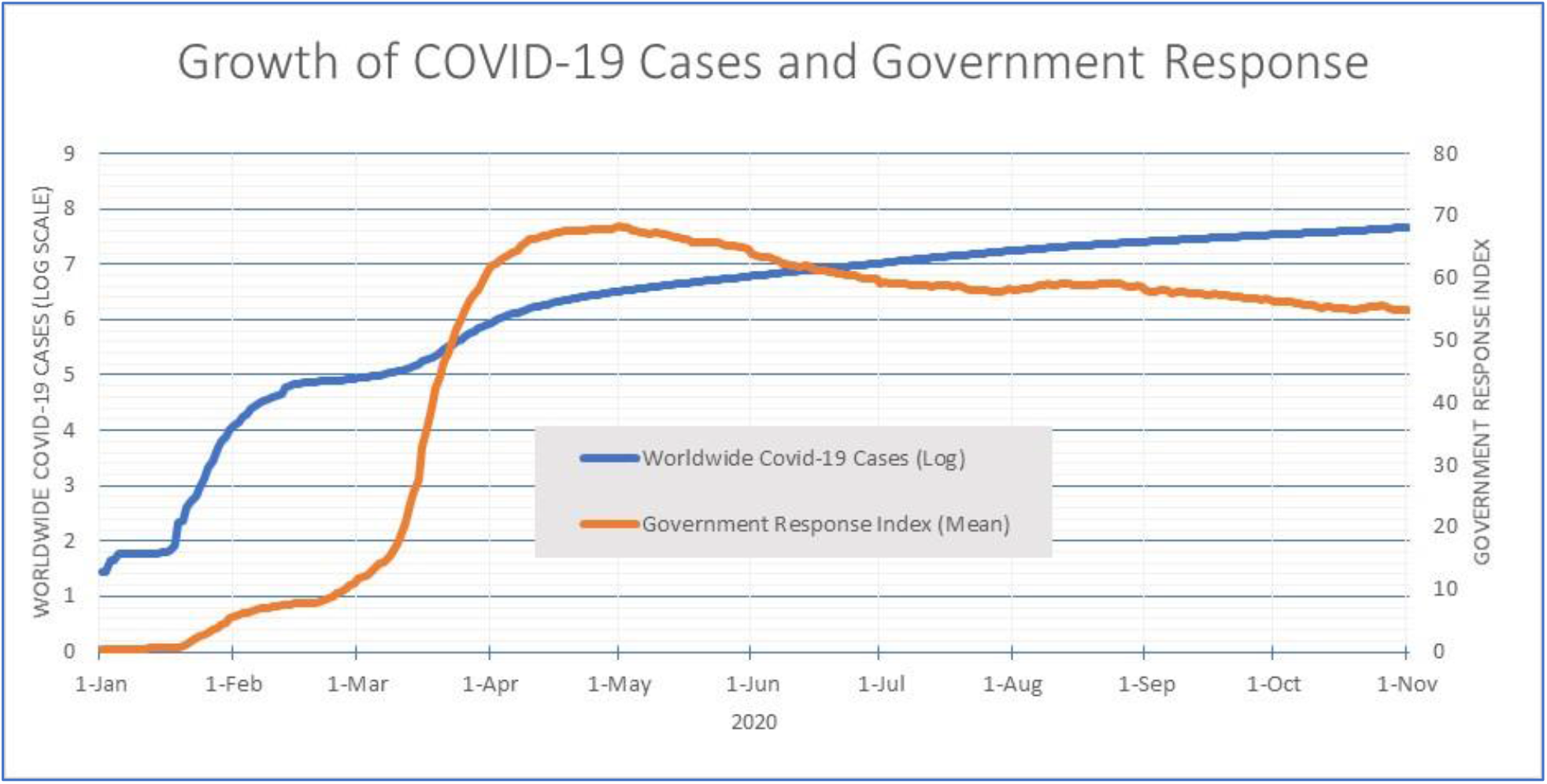
Calculated from Our World in Data, COVID-19 dataset (see footnote 4); and OxCGRT’s global mean index values for COVID-19 government response (see footnote 6).

Governments around the world have responded to the COVID-19 pandemic with a range of policy responses including workplace and school closings, requests and orders to stay at home, controls on domestic and international travel, financial and economic support, and multiple healthcare and public health measures.^5^ The Oxford COVID-19 Government Response Tracker (OxCGRT) indicates that some governments were undertaking policy responses in January 2020. However, the greatest surge worldwide in COVID-19 government responses occurred in March and April of 2020, continuing – with some fluctuations – throughout 2020, as measured by OxCGRT’s global mean index values for COVID-19 government response for over 180 countries (Figure 1).^6^

### Implications of the COVID-19 pandemic on scientific research

Published reports and articles indicate that the COVID-19 pandemic has had multiple consequences for scientific research.

For scientists working directly on COVID-19 research, including in epidemiology, virology, testing and diagnosis, therapies and treatments, medical equipment, vaccine development, and public health responses, it has been a period of urgent focus with expectations of rapid results. Worldwide, scientists have been activated to respond to COVID-19, using both existing resources and a massive injection of new R&D funds by research sponsors.^7,8^ Research sponsors have shortened proposal review times and accelerated award processes for new COVID-19 R&D funding.^9^ More than 175 new COVID-19 R&D programs and projects have been identified by OECD in in multiple locations (including in Europe, the US, Canada, Australia, China, Brazil, Russia and India).^10^ Another estimate suggests that, since January 2020, more than $9 billion in government, philanthropic, and industry funding has been committed globally for COVID-19 R&D.^11^ Much of this funding has gone to COVID-19 R&D by private companies, although universities and other public research organizations have also been supported, with the US being by far the largest sponsor, followed by Canada, Germany and the UK.^12^

The urgency of the pandemic and the extraordinary new R&D focus on COVID-19 has resulted in a massive increase in COVID-19 research publications. There has been a sharp growth in papers addressing COVID-19 topics, with US, Chinese, UK, and other European researchers anchoring the largest research networks.^13^ There has been growth not only in peer-reviewed journal articles but also in preprints (non-peer reviewed, publicly available papers) that address COVID-19 topics, with new tools and databases emerging to track and sort these papers.^14^ Cautions have been raised about the quality of COVID-19 “speed science,” especially among non-reviewed preprints.^15^ However, one early (April 2020) study found that more than three-quarters of the 11,686 COVID-19 papers identified at that point were journal papers (compared with preprint papers in repositories, many of which were pre-publication versions of papers in processes of journal submission).^16^ The study noted that the three largest open-access repositories for COVID-19 papers were PMC and PubMed, medRxiv and BioRxiv.

Meanwhile, particularly since March 2020 as governments expanded responses to curtail the spread of the pandemic, scientists in non-COVID-19 research domains have endured multiple disruptions to regular research work.^17,18,19^ Research facilities, laboratories, and labs have been closed for long periods in many countries. Across all fields, large numbers of researchers have spent months working from home, using remote access where possible. The recruitment of doctoral and early career researchers (especially internationally) has slowed, where it was not suspended.^20^ Research-related fieldwork and other travel has halted, while online video conferencing for team meetings, workshops and conferences has boomed. With school closures and social distancing, scientists with family and other caring responsibilities have faced added pressures, with effects on research productivity especially experienced by women.^21,22^

The closure of research facilities in 2020 resulted in benchwork, experiments and trials being stopped and the blocking of access to labs, instruments, and equipment. For example, more than 90% of labs at MIT were shut by March 20.^23^ The only exemptions were for labs working with coronavirus research, expensive materials, and certain animal experiments. As elsewhere, researchers shifted to a virtual environment, including using (and publishing with) already available data, or reorienting projects to involve more simulation work. In some instances, new lab automation was introduced. However, remote working for many months has presented challenges. For non-COVID 19 cancer research, access to labs and samples, the pausing of patient enrolment on clinical trials, and face-to-face interactions with colleagues have been major concerns.^24^ In contrast, for some other non-lab bound scientists, the worldwide slowdown or ceasing of human activities caused by the pandemic presented unique observational opportunities. New opportunities emerged to study impacts of reduced traffic, ocean shipping, pollution, noise, and other human activities on the environment, cities, and the natural world.^25,26^

Towards the middle and latter months of 2020, research labs began to re-open in a growing number of countries. In this phase of ramp-up, labs typically re-opened with capacity reductions or controlled shift work due to social distancing and other preventative measures.^27,2829^ At the UK’s Institute for Cancer Research, the proportion of time researchers spent in labs dropped from 53% pre-pandemic to 5% in lockdown, returning only to 34% many months later during phased access.^30^

The reopening of research labs for non-COVID-19 research has proceeded cautiously, even in locations, such as China, where the spread of COVID-19 has been controlled following the initial outbreak. In other countries, including in the UK and other parts of Europe, there have been phases of ramping-up of some research activities, followed by subsequent lockdowns in the Autumn of 2020.

The outlook for the large-scale rollout of safe and effective vaccines now appears optimistic for 2021. However, there remain uncertainties in many countries as to when labs can fully open, without social distancing, and when international travel will fully resume. Some work practices established in the pandemic may persist (including continued extensive use of virtual team working and virtual conferencing), while there are likely to be enduring impacts for many researchers in terms of lost experiments, reoriented topics, and career development.

### Identifying COVID-19 publications: A bibliometric search approach

To provide an update to bibliometric studies undertaken earlier in the pandemic, the paper probes trends in COVID-19 publications and non-COVID-19 publications as far as is possible in 2020.

Complete publication data for the full year of 2020 is not available at the time of this analysis. Access is available for publication datasets through to September to November of 2020, allowing about 9 to nearly 11 months of publication data to be analyzed (depending on the database). Two publication databases are used. The first is the PubMed database, maintained by the National Center for Biotechnology Information at the US National Library of Medicine (NLM). This allows searches of more than 31 million journal records and abstracts in biomedical, health and related fields, including more than 8 million full-text records.^31^ The second database is the Web of Science (WoS), maintained by Clarivate Analytics, which contains 78 million publication records from about 21,000 peer-reviewed journals in more than 250 fields of science, social science, and arts and humanities.^32^ Both PubMed and WoS have worldwide coverage, although they predominantly contain English-language records.^33^

To identify COVID-19 records in PubMed, the following COVID-19 bibliometric search approach developed by Search Technology and Georgia Tech was used:^34^

#### PubMed

“COVID-19”[All Fields] OR (“coronavirus”[MeSH Terms] OR “coronavirus”[All Fields]) OR “Corona virus”[All Fields] OR “2019-nCoV”[All Fields] OR “SARS-CoV”[All Fields] OR “MERS-CoV”[All Fields] OR “Severe Acute Respiratory Syndrome”[All Fields] OR “Middle East Respiratory Syndrome”[All Fields]

This search approach was applied to PubMed to obtain summary totals for COVID-19 records for all years. Additionally, for detailed analysis based on aggregations of individual records, the paper uses a publicly available database of COVID-19 PubMed records made available by Search Technology based on the above search approach. This database contained 56,463 COVID-19 records from PubMed published in 2020 through to September 22, 2020 (All documents).^35^

To identify COVID-19 records in WoS, the same bibliometric search approach was adapted for the WoS Advanced Search facility:

#### WoS

TS=(“COVID-19” OR Coronavirus OR “Corona virus” OR “2019-nCoV” OR “SARS-CoV” OR “MERS-CoV” OR “Severe Acute Respiratory Syndrome” OR “Middle East Respiratory Syndrome”)

This search approach was applied to the WoS (Indexes=SCI-EXPANDED, SSCI, A&HCI, CPCI-S, CPCI-SSH, ESCI) refined by document types (article, early access, review, or proceedings paper). After capturing records and cleaning, this resulted in 44,022 COVID-19 WoS records, published 1969 through to October 24, 2020. Aggregated analysis was also undertaken for all WoS publication records (with same Indexes and document types, for 2020) downloading the summary results (but not individual records).

VantagePoint desktop text mining software was used for further processing and analysis of the two datasets containing the WoS and PubMed records.^36^

The next parts of the paper present analyses using the WoS and PubMed publication data. In interpreting these analyses, keep in mind the differences between the two databases. As noted, the WoS covers all fields of science, social science, and arts and humanities. The analysis that follows includes articles, early access, review, or proceedings papers from reviewed sources. Articles and reviews comprise about 95% of the COVID-19 WoS dataset. Preprints and chapters in edited volumes are not included, and there is some lag (most relevant for 2020 data) in review and publication processes. PubMed is focused on biomedical, health and related fields, but there is a wider sweep in the records reported including Medline (containing publication records from evaluated journals), papers from other journals, chapters and preprints (the latter with less time lag in posting).

### Growth of COVID-19 publications in 2020

Research related to various forms of coronavirus has been undertaken for several decades, with publication at a relatively low level, e.g. at around 150-200 annual publications in the late 1990s and early 2000s.^37^ There was a spike in research publications with about 700 WoS and 4,000 PubMed publications over the 3-year 2003-2005 period, with the global emergence of severe acute respiratory syndrome (SARS), a distinct coronavirus (SARS-CoV). Subsequently, coronavirus-related publications continued at annual levels of around 650 and 760 in the WoS and PubMed in the 2006 to 2019 period.

The publication profile, of course, changed dramatically in 2020, with the rapid worldwide spread of COVID-19. The urgent focus of scientific attention generated by the pandemic, as noted earlier in this paper, has been associated with a massive increase in publication outputs on COVID-19 topics. From January to November 2020, more than 38,000 WoS and 78,000 PubMed publications were recorded. Reflecting the upward step in the volume of COVID-19 publications is a noticeable shift in 2020 in the proportion of all scientific publication efforts devoted to pandemic-related research. When compared with 2019, the share of COVID-19 publications in 2020 in all publications rose from 0.03% to 1.8% in the WoS and from 0.07% to 5.3% in PubMed (Figure 2).

**Figure 2.**
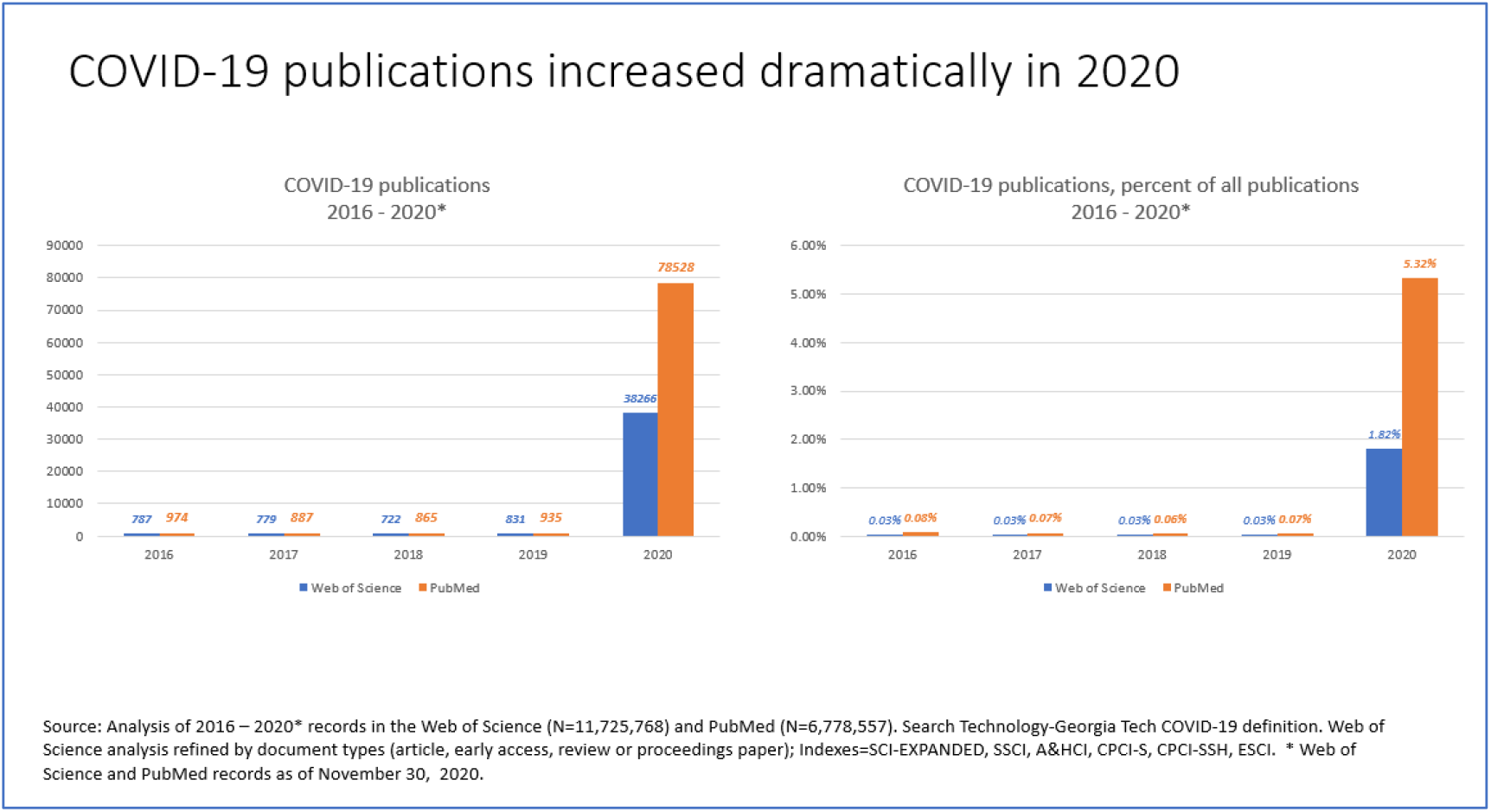

On a month-by-month basis in 2020, COVID-19 research publications in PubMed began to accelerate in February and March 2020 (when the global extent of the pandemic was fully realized, and government responses heightened). There was a further increase in rate of PubMed COVID-19 outputs from April through to the Summer of 2020. (Figure 3.) This analysis is based on PUBMED records published in 2020 as of September 22, 2020. Publication tallies for more recent months are not completely captured. In other words, the apparent downward PubMed profile from August 2020 is based on incomplete record capture. More publications will be added in the more recent months when 2020 is more fully indexed (by the first part of 2021). WoS COVID-19 publications show comparable trends, except with a lower rate of take-off (than for PubMed publications) in Spring 2020. Again, the WoS dataset (containing COVID-19 publication records as of October 24, 2020) is not complete for more recent months.

**Figure 3.**
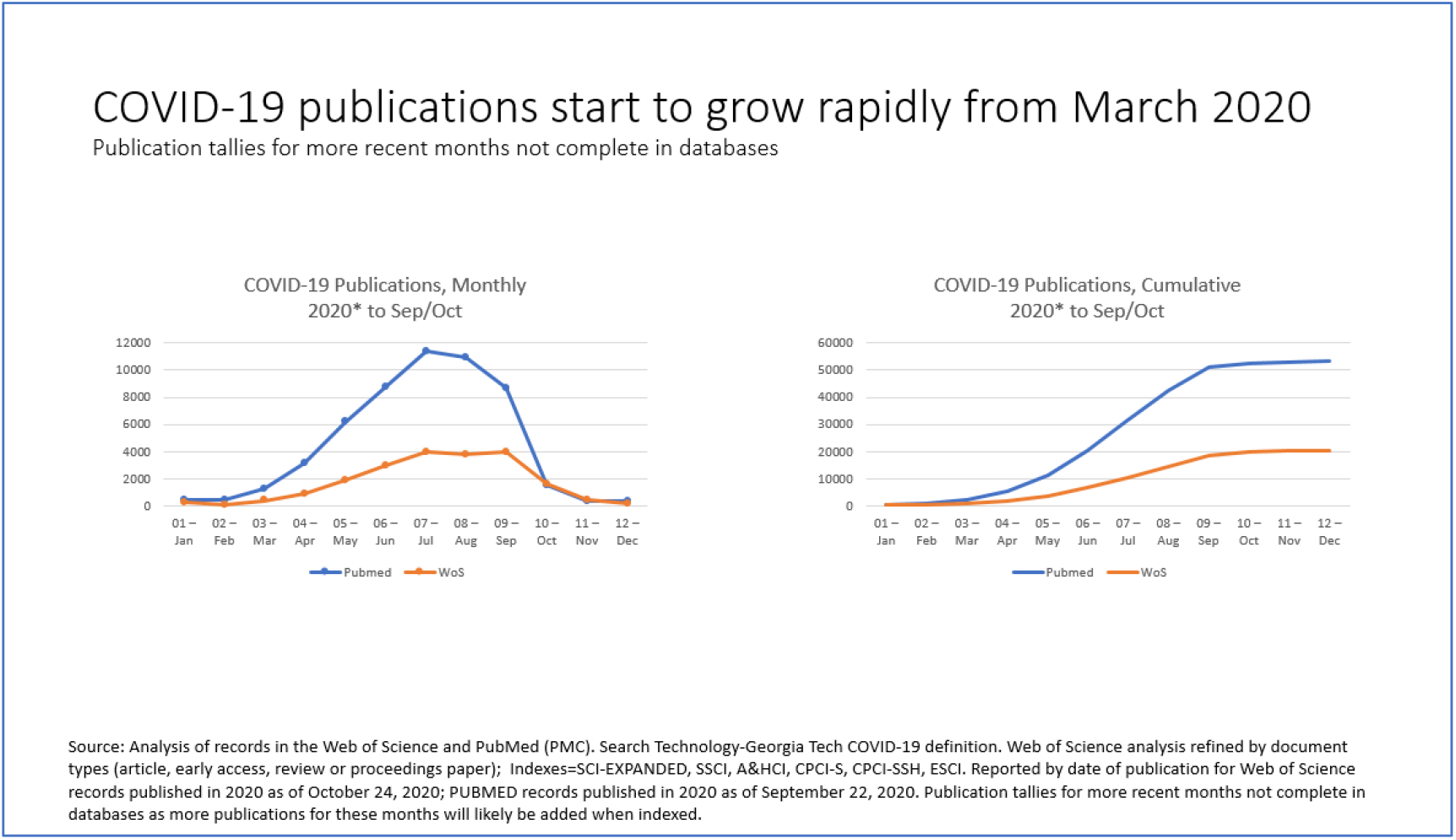

### COVID-19 publications in 2020 by countries and organizations

By country, based on author affiliation location, for both WoS and PubMed COVID-19 research publications in 2020, the leading producer is the US, followed by China, the UK, and Italy (Figure 4). The mid-size developed (OECD)^38^ countries of France, Canada, Spain, Germany, and Australia comprise all but one among other locations in the top ten. India is positioned fifth by number of 2020 COVID-19 papers, with two other non-OECD countries – Brazil and Iran – placed eleventh and twelfth.

**Figure 4.**
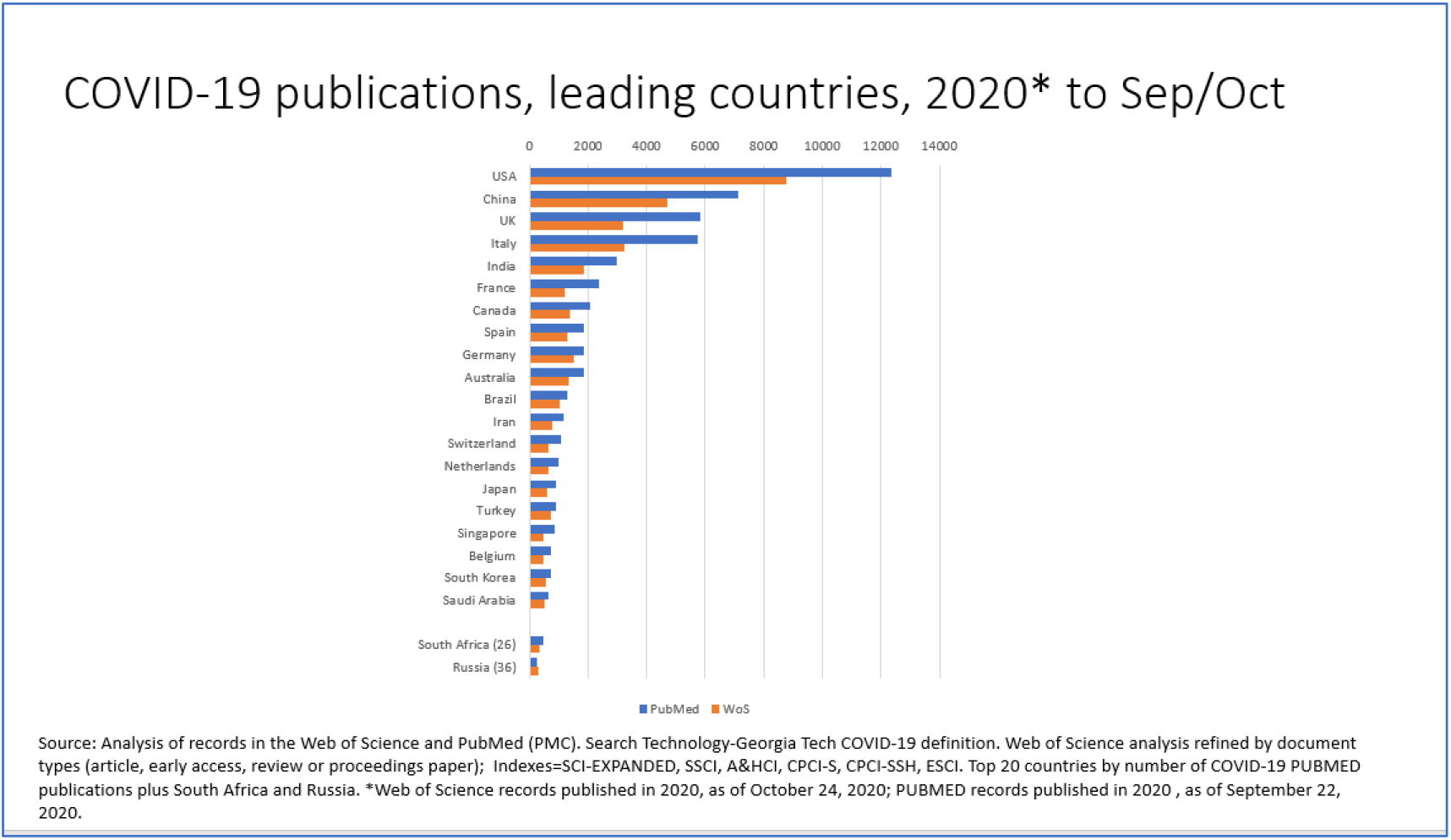

Based on WoS publications, the positioning of the US and China is reversed when publications across all fields is considered. The US contributed 29.1% of 2020 publications in the COVID-19 WoS dataset, compared with 22.3% of all WoS publications over the period January 2020 through to November 2020, a ratio of Covid-19 to all papers of 1.30.^39^ The comparable numbers for China are 15.6% of COVID-19 WoS papers and 23.6% of all WoS papers, with a ratio of Covid-19 to all papers of 0.66. Understanding the reasons for this difference needs further investigation, but it probably reflects the strong and relatively larger base of biomedical research in the US and the influence of rapidly available added COVID-19 research funding. Additionally, it is important to note that China has published and made available a large volume of WoS (as well as PubMed) papers on COVID-19.

Among other countries, the UK, Italy, India and Canada published in 2020 (to date) a relatively larger share of COVID-19 WoS papers when compared with all WoS papers (ratios of 1.46, 2.43, 1.15, and 1.15 respectively). France and Germany are among countries publishing COVID-19 papers in 2020 at comparable or relatively lower rates when compared with all WoS papers (1.03 and 0.83 respectively). Authors affiliated with institutions in Russia published COVID-19 WoS papers at relatively lower rates when compared with all Russian WoS papers (ratio of 0.42).

Researchers from a wide range of organizations (based on author affiliations) in multiple countries have been contributing to the rapid expansion of COVID-19 research papers in 2020. Of the top 20 organizations by PubMed COVID-19 publications in 2020 (to September 22), nine are based in the US, four in China, two in the UK, and one each in Singapore, Italy, India, Canada, and Iran (Figure 5). Mostly, the leading organizations are research universities and specialized medical research institutions, although there is notable representation from identifiable hospitals and clinics.^40^ For WoS COVID-19 publications in 2020 (to October 24), organizations from fewer countries are represented in the top 20, but the top part of that list is comparable, led by eight organizations based in the US, six in China, and three in the UK (Figure 6).

**Figure 5.**
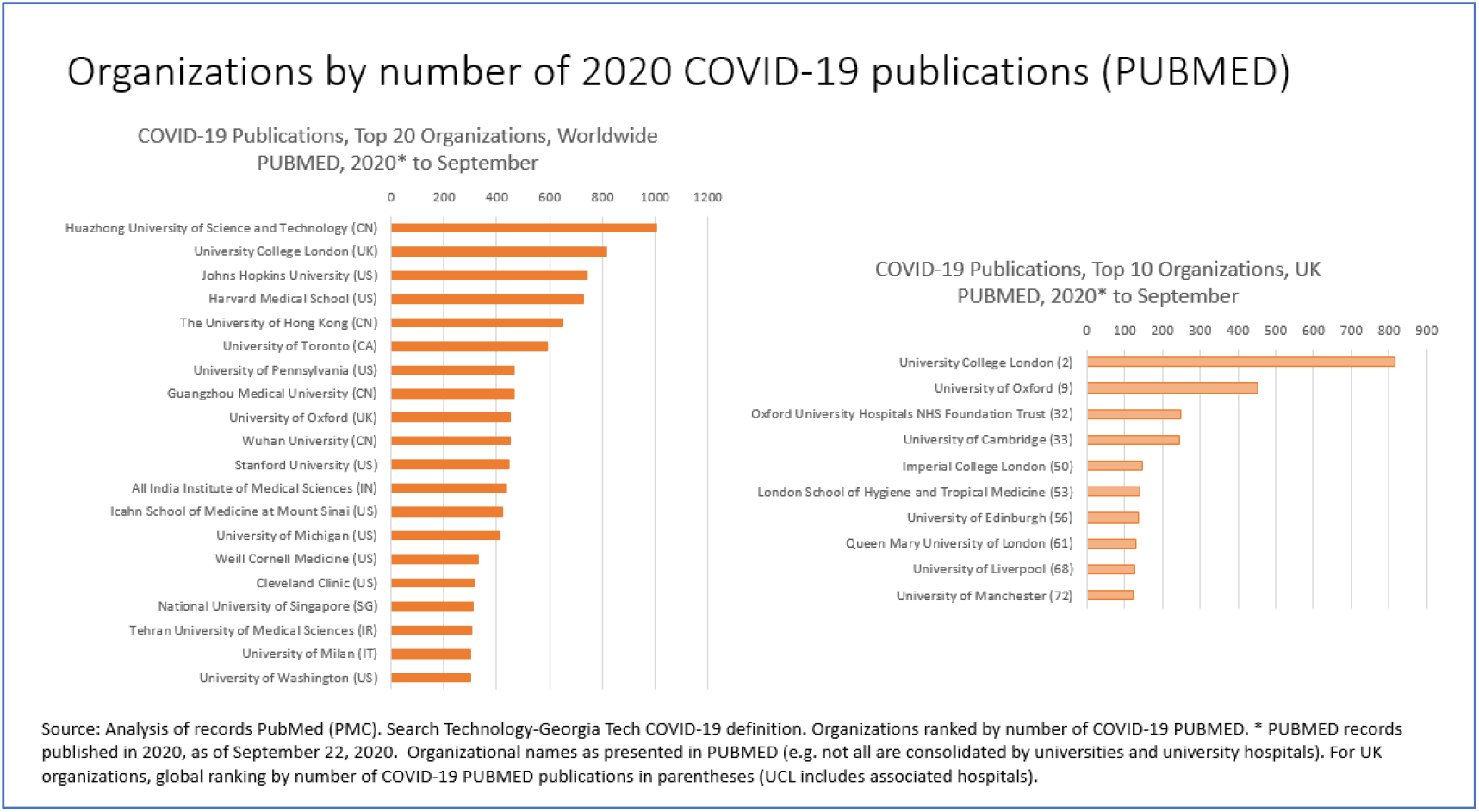

**Figure 6.**
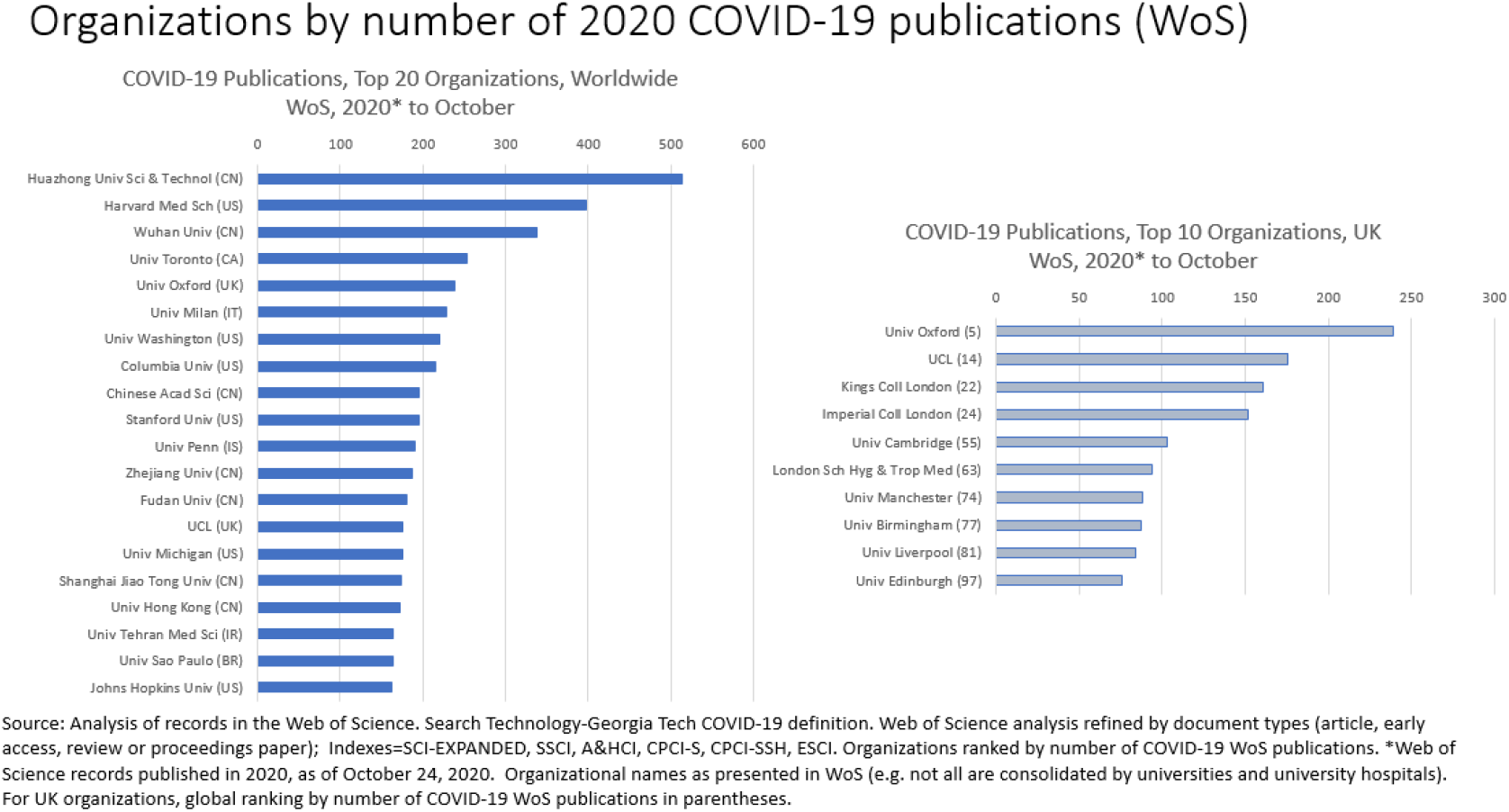

Huazhong University of Science and Technology (in Wuhan, Hubei province, China) is the leading publisher of both PubMed and WoS COVID-19 papers, by volume of papers. Wuhan University is tenth organization by PubMed COVID-19 papers and third by WoS COVID-19 papers, again by volume of output.^41^ The University of Hong Kong is also a leading producer of PubMed COVID-19 outputs. Johns Hopkins University, the Harvard Medical School, the University of Toronto, and the University of Pennsylvania in North America, and University College London and Oxford University in the UK, are leading producers of PubMed COVID-19 publications. These are all large research institutions with significant medical and public health research capabilities. By output of WoS COVID-19 papers, research institutions in China, North America, and Europe (the UK and Italy) comprise the top ten.^42^

### COVID-19 Publications by Subject Categories

The journals included in the WoS publication database are classified by over 250 “subject categories.”^43^ These subject categories (SCs) are broadly equivalent to disciplines, but this is not a perfect match.^44^ Journals can and do cover multiple fields of science. In the WoS scheme, journals (and the papers they publish) may be classified in up to 6 SCs, although about 60% of journals in WoS are classified in only one SC, with another 30% classified in two SCs.^45^ Some journals are explicitly interdisciplinary. Reflecting this there are nine WoS SCs that are designated as multidisciplinary, including “Multidisciplinary Sciences” (which includes such journals as *Nature* and *Science* that publish across all fields of science).

The journal SC (or combination of SCs) is allocated to every paper in the journal, although individual papers within a particular journal may be varied in their subject matter. Additionally, as scientific fields evolve and novel fields form, new journals typically emerge. This presents classification challenges and lags, although SCs and constituent journals are periodically updated to reflect changes in fields, new fields, and repositioning of journals. While it is convenient and common to use WoS SCs to overview the distribution of published outputs by various scientific fields, these caveats about the classification scheme should be kept in mind.

To provide a basis over time for comparison, WoS COVID-19 publications are analyzed for two time periods: 2017-2019^46^ and 2020 (through to October 24). (Figure 7.) In the earlier period, “Virology” was the leading SC category, with 562 coronavirus WoS publications, or 24.4%) of this SC’s total from 2017 to 2019 (three complete years). In this earlier period, the second and third SCs by 562 coronavirus WoS publications were “Infectious Diseases” (354 publications) and “Microbiology” (285 publications).

**Figure 7.**
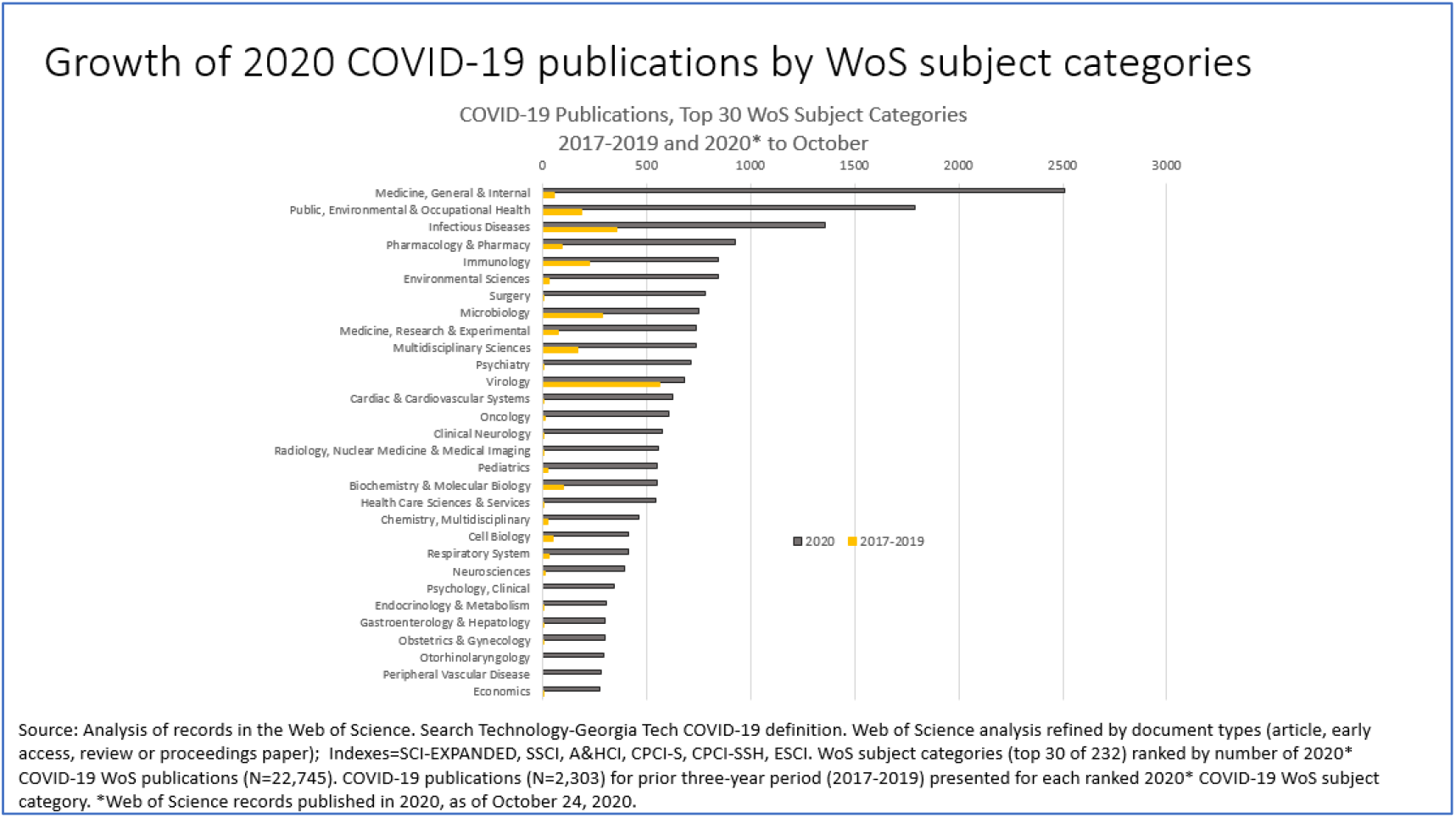

Together, more than half (52.1%)^47^ of earlier period coronavirus outputs were classified in these three SCs, reflecting a research focus on the properties and microbiology of the virus itself.

In 2020, the profile of COVID-19 research publications dramatically changed – with an upsurge in the volume of publications and engagement of scientists from a wider array of research fields. There was a massive increase in “Medicine, General & Internal” (over 2,500 WoS COVID-19 publications in 2020 to date, compared with just over 50 WoS coronavirus publications in the prior three-year period). Along with large increases of 2020 COVID-19 WoS publications in the SCs of “Public, Environmental & Occupational Health,” “Pharmacology & Pharmacy,” “Environmental Sciences,” “Surgery,” and “Medicine, Research and Experimental,” this represents an understandable and rapid broadening of COVID-19 research activities into the domains of treatment, therapy, contagion, and public health. Multiple other medical fields with little prior coronavirus research activity saw significant increases in 2020 COVID-19 WoS publications, including “Psychiatry,” “Cardiac &, Cardiovascular Systems,” “Oncology,” “Clinical Neurology,” “Radiology,” “Nuclear Medicine & Medical Imaging,” and “Pediatrics” (see Figure 7). The SCs of “Infectious Diseases,” “Microbiology,” and “Virology,” as with “Immunology”, “Biochemistry & Molecular Biology,” and “Cell Biology”, all saw surges in 2020 COVID-19 WoS publications as researchers further analyzed the nature and pathology of the virus and explored human impacts and responses. Research on vaccines is categorized across several SCs: the leading categories for COVID-19 WoS publications related to vaccines are “Immunology,” “Virology”, “Medicine, Research and Experimental,” “Microbiology,” and Biotechnology & Applied Microbiology.”

### COVID-19 Pivots in 2020 by Research Fields

With the urgency and global spread of the pandemic in 2020, scientists around the world have substantially expanded their COVID-19 research efforts. One outcome of this, as we have seen, is a massive increase in the volume of PubMed and WoS publications on COVID-19 topics. It is evident, from the analysis in the previous section of WoS subject categories, that researchers in medicine, biological sciences, and public health domains have greatly expanded COVID-19 publication outputs in 2020. The entry of “Economics” (at 30^th^ place) in the top 30 SCs by number of WoS COVID-19 publications in 2020 (Figure 7) highlights COVID-19 research activity in the social sciences. This is comprehensible: while the global pandemic has caused tragic public health consequences and massive impacts on health care systems, it has resulted in huge effects worldwide across all aspects of life and human activity, economy, society, community, and government.

Given the scale and scope of the pandemic, scientists across a wide range of research fields will surely be interested in, and concerned about, the impacts and consequences of COVID-19 in its many and varied forms. Those interests and concerns may take different forms including policy and community engagement as well as reoriented or new research projects related to COVID-19 topics. Publication opportunities from these activities and projects likely varies across fields. As discussed earlier in this paper, large numbers of scientists not working directly on urgent COVID-19 medical, biological science, or public health topics have spent much of 2020 working from home or at reduced levels of access to their labs, field sites, or research subjects. This surely influences research paper productivity. In certain circumstances, output may increase, for example for scientists working directly on urgent COVID-19 topics or who have access to data that can be written up. We have clearly seen a massive increase in PubMed and WoS publications related to COVID-19. To an extent, this wave of new COVID-19 publications may have substituted for what would have been other outputs; or the urgency of the pandemic and the massive efforts of researchers (and perhaps a strong desire by some to be first to publish) may have generated increased publication productivity. In many other cases, there may be adverse effects on publication productivity, where research has slowed or paused or where researchers have been preoccupied with family or caring responsibilities, online teaching (where researchers also teach), crisis management, or suffered personal or family health impacts.

It is important for many reasons (including research management, researcher career development, and policy learning) to assess how research productivity, as measured by publication outputs, has been affected during the pandemic, with comparison to pre-pandemic norms (and, of course, the post-pandemic era, when we reach that point). Such an analysis should account for field and institutional effects as well as potential differential effects by researcher gender, seniority, and roles. Any immediate (2020) analysis should also account for the significant variations in time lags to publication by journals in different fields^48^ and the impact of the pandemic on journal peer review processes.^49^

However, 2020 is not yet complete (at the time of analysis and writing). Indeed, a complete record of publications for the full year of 2020 will only be available only in 2021, as some time needs to elapse before all prior period publications are recorded in indexes such as PubMed or WoS. Hence, it is not yet possible to undertake disaggregated and controlled measures of research publication output by quantity for the full year of 2020 compared with 2019 (or, indeed, for earlier years).

These caveats noted, it is nonetheless possible to assess COVID-19 research pivots not only for medical, biological science, or public health fields, but across *all* domains of science, social science, and the humanities. A COVID-19 research pivot is defined as the extent to which the proportion of output in a particular research field has shifted to a focus on COVID-19 topics in 2020 (to date) compared with 2019. COVID-19 research pivoting is calculated by measuring publication outputs on COVID-19 topics in a field as a percentage of all publication outputs in that field for 2019 and 2020, then subtracting the 2019 percentage from the 2020 percentage. This results in a COVID-19 “pivot point” index. This is a simple method, but it is effective in normalizing outputs for the yet not quite complete year of 2020. There are some fields (for example, “Virology”) with a proportion of publication outputs captured by the COVID-19 bibliometric search term in 2019; the pivot point measure highlights the added extent to which the proportion of 2020 COVID-19 publication outputs increased over the prior year. Most research fields, however, had zero 2019 COVID-19 publication outputs; in this case, the pivot point measures the new extent to which research outputs in that field shifted to a COVID-19 focus in 2020, if at all.

The method is operationalized by applying the COVID-19 bibliometric search term to the full WoS online database as of October 24, 2020 (Indexes=SCI-EXPANDED, SSCI, A&HCI, CPCI-S, CPCI-SSH, ESCI) refined by document types (article, early access, review, or proceedings paper). For 2019, the total number of WoS records is 2,567,047, of which 828 are identified as COVID-19 publications. For 2020 (through to October 24), the total number of WoS records is 1,952,201, of which 30,143 are identified as COVID-19 publications. WoS subject categories (SCs) are used as a proxy for research fields. All 254 WoS SCs (as of October 2, 2020) are included in the analysis. (See Appendix for data table.)

Recall that not all 2020 publications are recorded yet (whether for papers with COVID-19 or non-COVID-19 topics). Whether a WoS publication count of about 1.9 million papers by the third week of October 2020 (or 2.1 million by the end of November 2020^50^) is significantly different from the comparable numbers at the same time in 2019 cannot be determined as that data is not available to the researcher (although that information would be available internally to the producer of the database). However, it can be observed that there has been an overall increase in the share of COVID-19 outputs in all WoS publications, increasing from 0.03% in 2019 to 1.5% in 2020 (to October 24).^51,52^ This represents an absolute and significant increase of about 29,300 COVID-19 papers in 2020 (to October 24). Most published papers recorded in the WoS in 2020 are *not* COVID-19 related (i.e. the whole of science has not pivoted to COVID-19 research). Nonetheless, by specific SCs, there are substantial variations, with some fields (as defined by WoS subject categories) making noticeable COVID-19 research pivots, while in most fields there is a small or non-detectable COVID-19 research pivot.

To analyze the extent of the relative shift into COVID-19 research topics by individual scientific fields in 2020 (to October 24), the COVID-19 research pivot (*RP*) measure is calculated for each WoS SC. For the 254 SCs, key *RP* values are: maximum = 15.8; mean = 1.5; median = 0.6; and minimum = 0.0. An asymmetrical and skewed distribution is indicated, with a long right tail of low to zero *RP* values. To visualize the distribution, the ranked *RP* measures by SC are presented by quintiles. The top quintile is named as comprising “high to medium” pivots, the second and third quintiles as “medium to low” pivots, and the fourth and fifth quintiles as “low to zero” pivots. Percentages of SC COVID-19 related outputs are denoted as *cPC19* and *cPC20* for 2019 and 2020 (to October 24) respectively.

In the first quintile, the SC with the highest COVID-19 research pivot is “Virology” (*RP*=15.8; *cPC19*=3.0%; *cPC20*=18.8%). (Figure 8.) The second highest is “Medical Ethics” (*RP*=15.0, *cPC19*=0.0%; *cPC20*=15.0%). Rounding out the top ten SCs with highest relative COVID-19 research pivots are “Infectious Diseases” (*RP*=9.5), “Medicine General Internal” (*RP*=7.0), “Otorhinolaryngology” (*RP*=6.4), “Public Administration” (*RP*=6.3), “Critical Care Medicine” (*RP*=5.6), “Public Environmental & Occupational Health” (*RP*=5.5), “Emergency Medicine” (*RP*=5.3), and “Health Care Sciences Services” (*RP=*5.2). For only three of these ten SCs is *cPC19* > 0.0, indicating that most of these fields have made a new and rapid research pivot into COVID-19 in 2020. Additionally, while six of the top ten SCs with the highest *RP* rankings are in medical specialties (including emergency and critical care) and biological sciences, the other four are in fields of medical ethics, public administration, public health, and health care delivery, highlighting the wide-ranging implications of the pandemic taken up by researchers.

**Figure 8.**
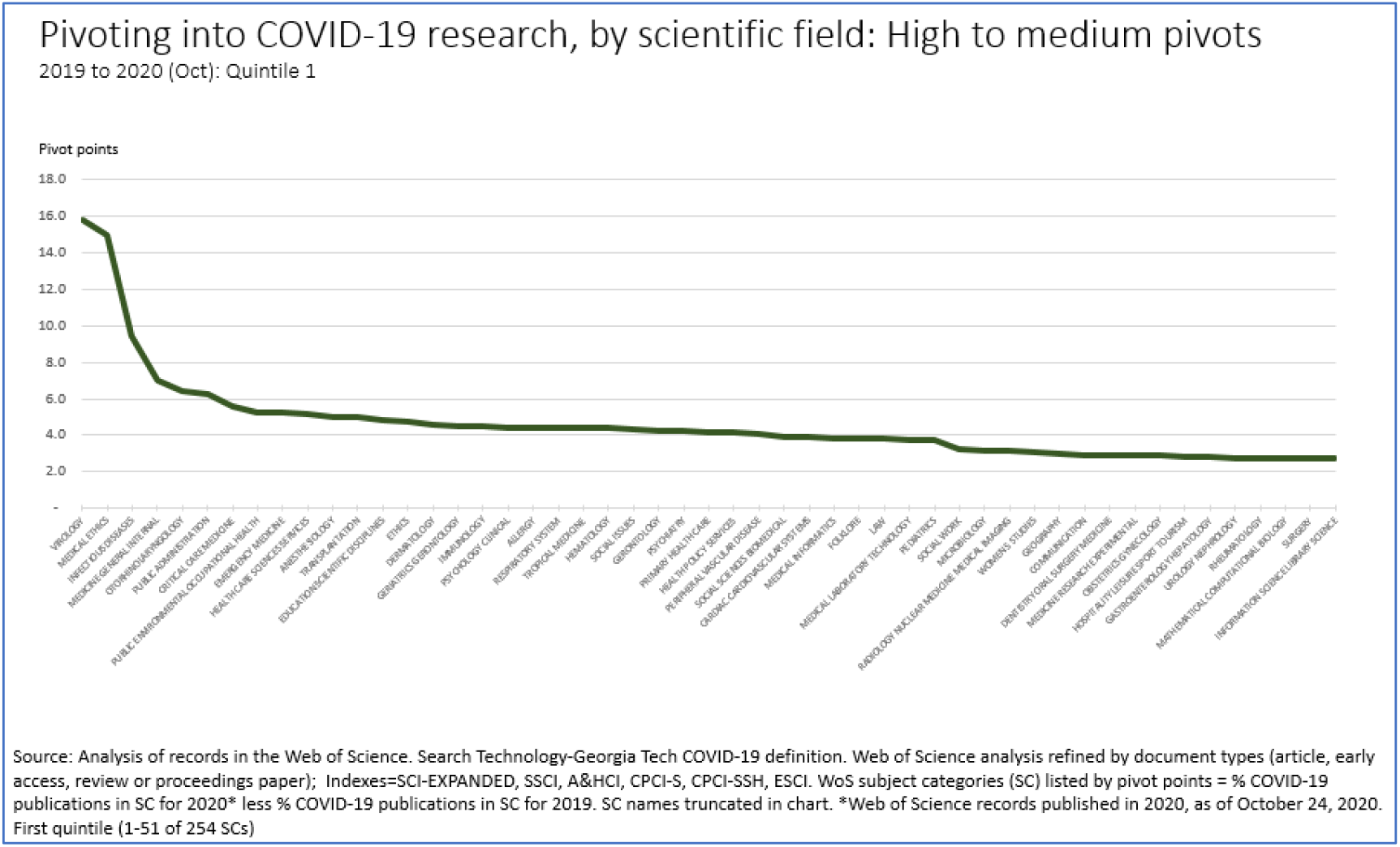

This pattern is replicated in the balance of the first quintile of high to medium *RPs*, with multiple medical specialties, biological sciences, and public health fields represented, alongside “Ethics,” “Social issues,” “Health Policy Sciences,” “Social Sciences, Biomedical,” and “Social Work.” Other entries in the medium *RP* sections of the first quintile also include “Folklore”, “Law”, and “Women’s Studies.”^53^Of the 51 SCs in the first quintile, 35 (69%) are indexed in the WOS SCI, 15 (29%) in SSCI, and 1 (2%) in A&HCI.^54^ In the first quintile, 2.7 < *RP* < 15.8 with a mean 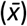 of 4.6.

The second and third quintiles (“medium to low” COVID-19 research pivots) also contain a mix of science, social science, and arts and humanities SCs. (Figure 9.) “Parasitology” (*RP*=2.7, *cPC19*=0.2%; *cPC20*=2.7%) and “Clinical Neurology” (*RP*=2.6, *cPC19*=0.0%; *cPC20*=2.6%) lead this intermediate set of quintiles, ranked 52 and 53 (of 254) respectively. The entries from SSCI and A&HCI in this set are led by “Area Studies” (*RP*=2.6, *cPC19*=0.0%; *cPC20*=2.6%) and “Cultural Studies” (*RP*=2.0, *cPC19*=0.0%; *cPC20*=2.0%). Although “Economics” is represented in 30^th^ position by absolute number of COVID-19 publications (Figure 7), it is more moderately ranked (89 of 254) in terms of its relative COVID-19 research pivot (*RP*=1.4, *cPC19*=0.0%; *cPC20*=1.4%). Economics is one of the largest producers of SSCI WoS publications; however, even though 350 COVID-19 related papers were identified in 2020 (to October 24), this was a relatively small proportion of its total SC output (25,325) to the search date.

**Figure 9.**
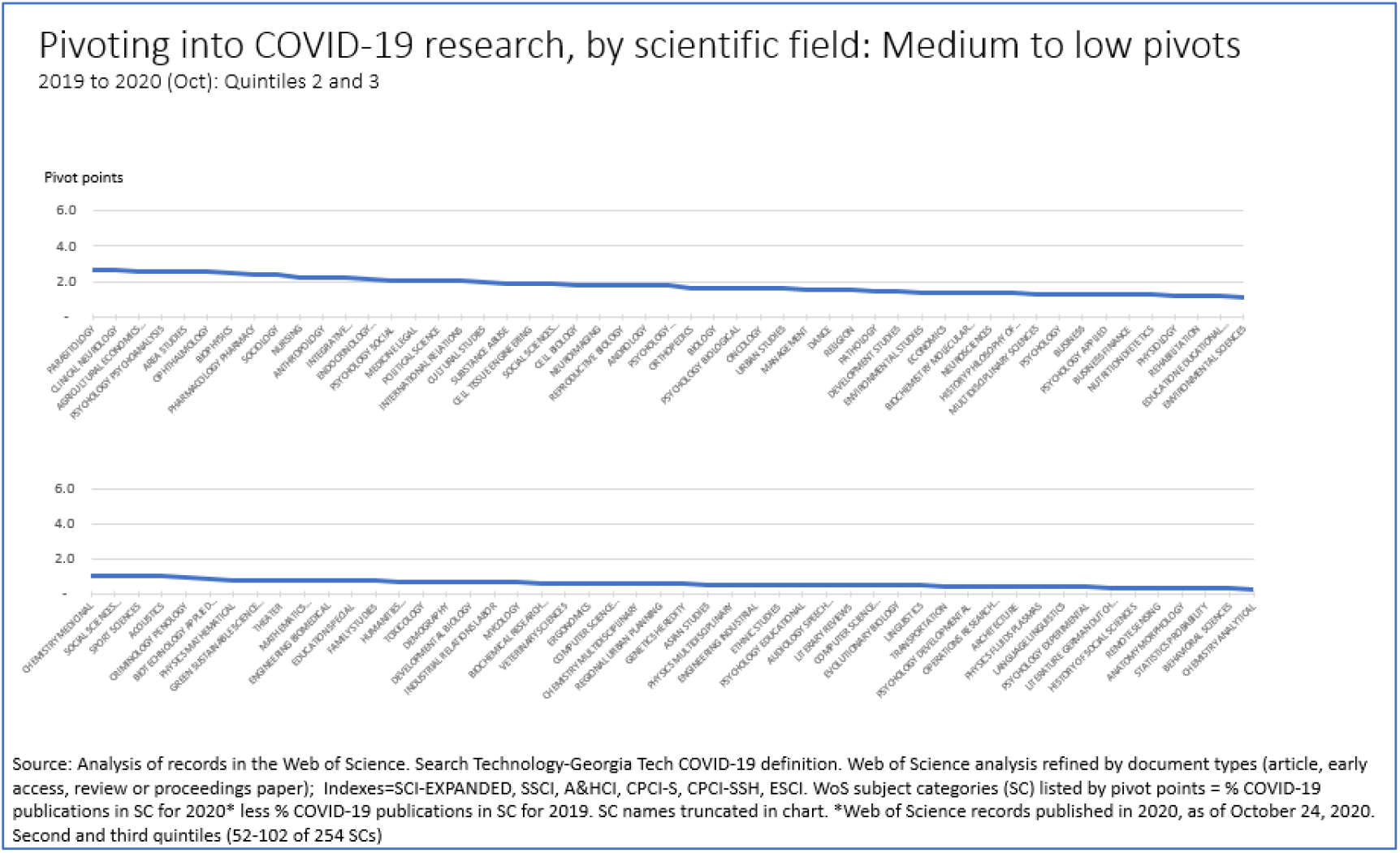

Of the 101 SCs in the second and third quintiles, 55 (54%) are indexed in SCI, 34 (34%) in SSCI, and 12 (12%) in A&HCI. In this medium to low set, 0.3 < *RP* < 2.7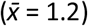. In the last set, the fourth and fifth quartiles, there are 102 SCs, of which 76 (75%) are indexed in SCI, 7 (7%) in SSCI, and 19 (19%) in A&CHI. In this “low-to zero” set, 0.0 ≤ *RP* < 0.3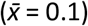. In 37 SCs in the lowest quintile, no COVID-19 related papers were identified out of more than 216,000 WoS publications for 2020 (to October 24). (Figure 10.) Again, note that publication data for 2020 is incomplete; researchers in these SCs may have preprints, papers or forthcoming and other COVID-19 research contributions not captured by this analysis.

**Figure 10.**
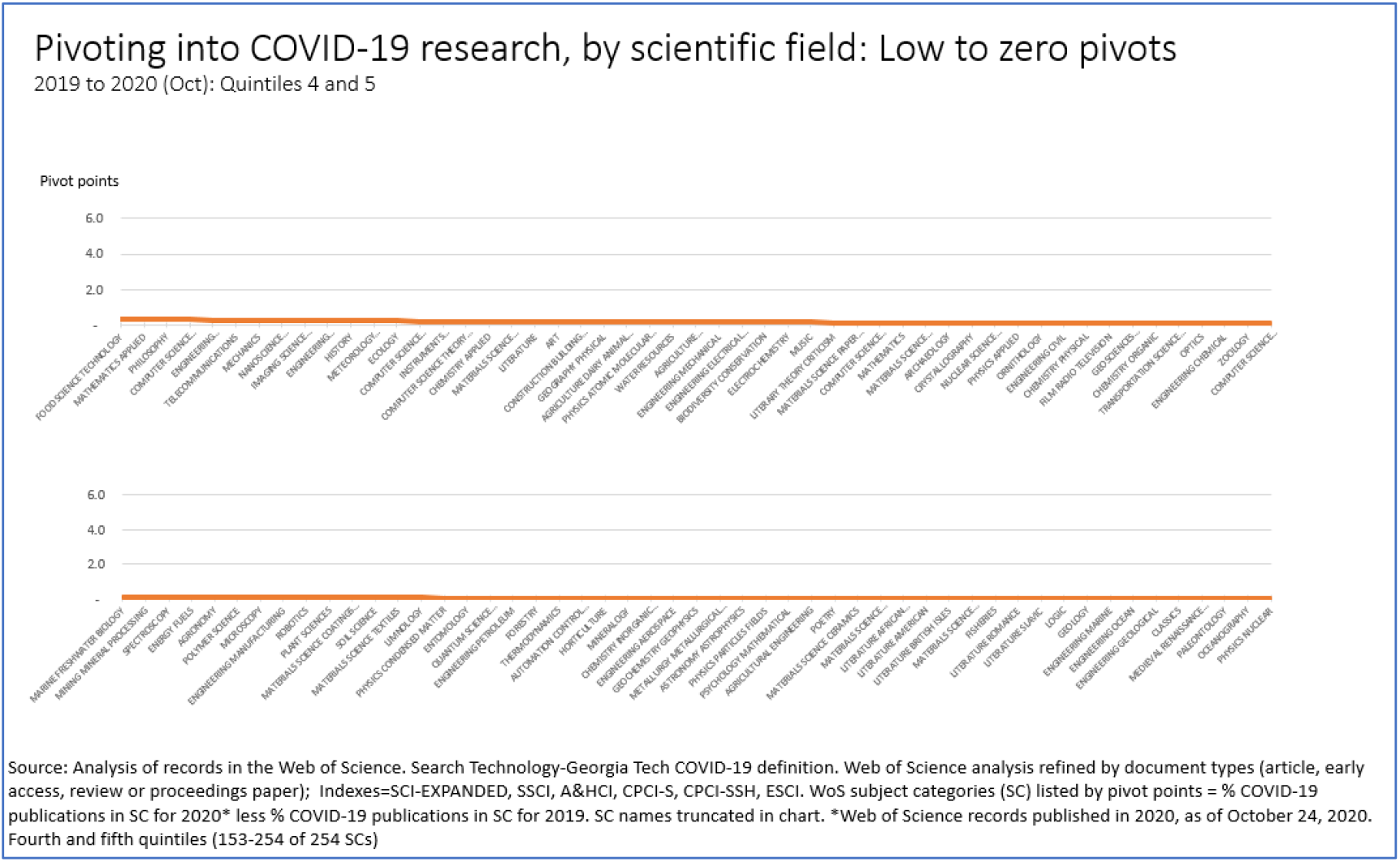

### Concluding Comments

The analyses presented in this paper should be regarded as preliminary. In 2020, scientific research around the world has been massively impacted by the global COVID-19 pandemic; these impacts are still in progress and will likely continue well into 2021. The full consequences and implications of COVID-19 and the pandemic on science, researchers and research institutions will take some further time, possibly years, to be wholly comprehended.

The paper has used available secondary and bibliometric data to analyze one aspect of the pandemic’s impacts on research, focusing on publication outputs (through to the September to November of 2020, depending on the databases used). Large-scale publication data is available in the PubMed and WoS database, although complete publication records for 2020 will not be available until the early parts of 2021. This caveat should be kept in mind, along with other limitations in using publication outputs to assess research performance, differences in content between PubMed and the WoS, and variations by fields and countries in approaches to publishing in sources that are recorded in these two publication databases.

The analysis finds a massive absolute increase in PubMed and Web of Science papers directly focused on COVID-19 topics, especially in medical, biological science, and public health fields, although this is still a relatively small proportion of publication outputs across all fields of science. Using Web of Science publication data, the paper examined the extent to which researchers across all fields of science have pivoted their research outputs in 2020 to focus on topics related to COVID-19. Significant variations are found by specific fields (identified by Web of Science Subject Categories). In a top quintile of fields – not only in medical specialties, biomedical sciences, and public health but also in subjects in social sciences and arts and humanities – there are relatively high to medium research pivots. In lower quintiles, including other subjects in science, social science, and arts and humanities, low to zero COVID-19 research pivoting is identified. It was noted that researchers may have preprints, papers or forthcoming and other COVID-19 research contributions not captured by the WoS analysis.

## Appendix 1

Data table of WOS publications used to calculate 2020 (to October 24) COVID-19 research pivots. See next pages. Explanation of headings and data sources is at the end of the table.

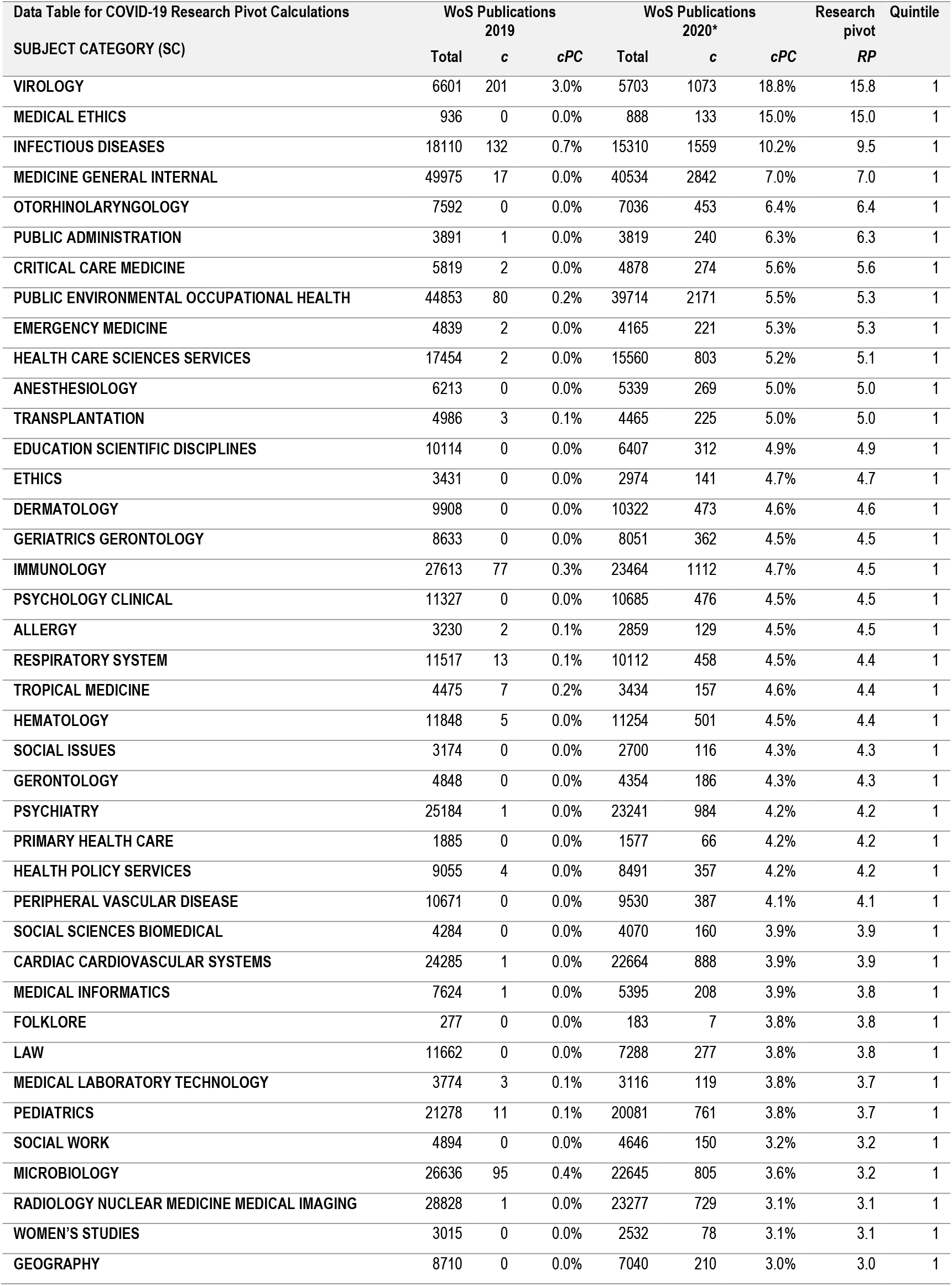
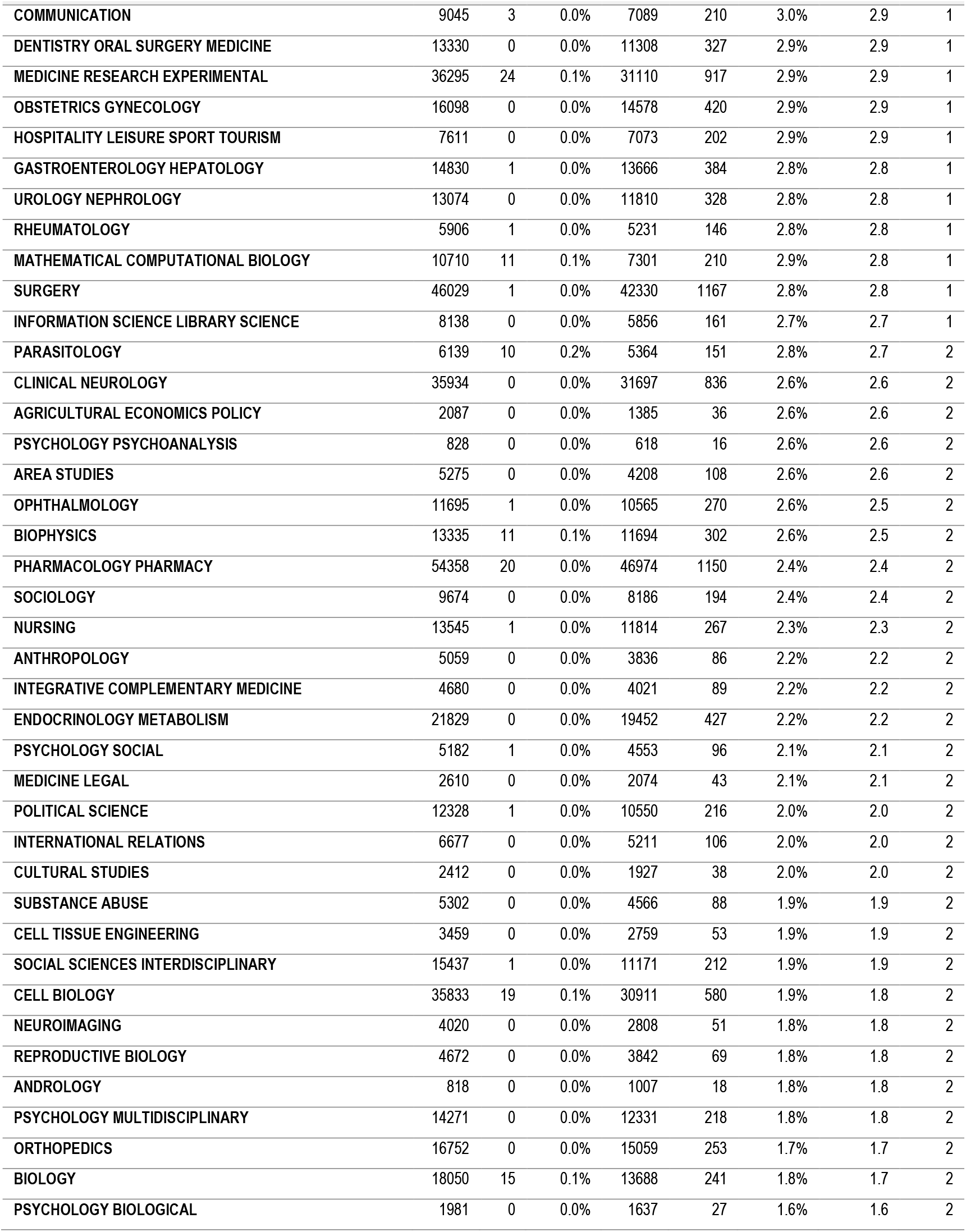
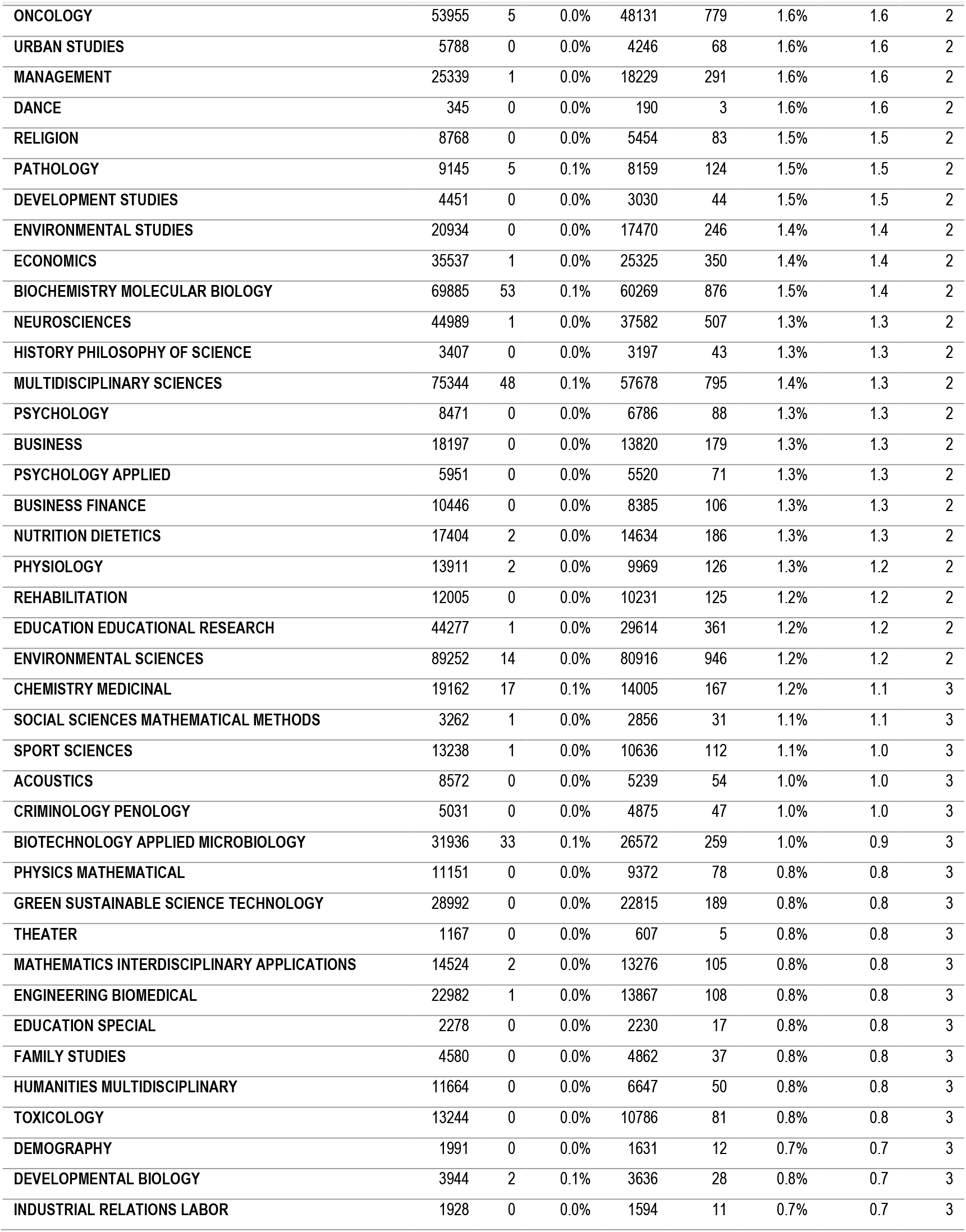
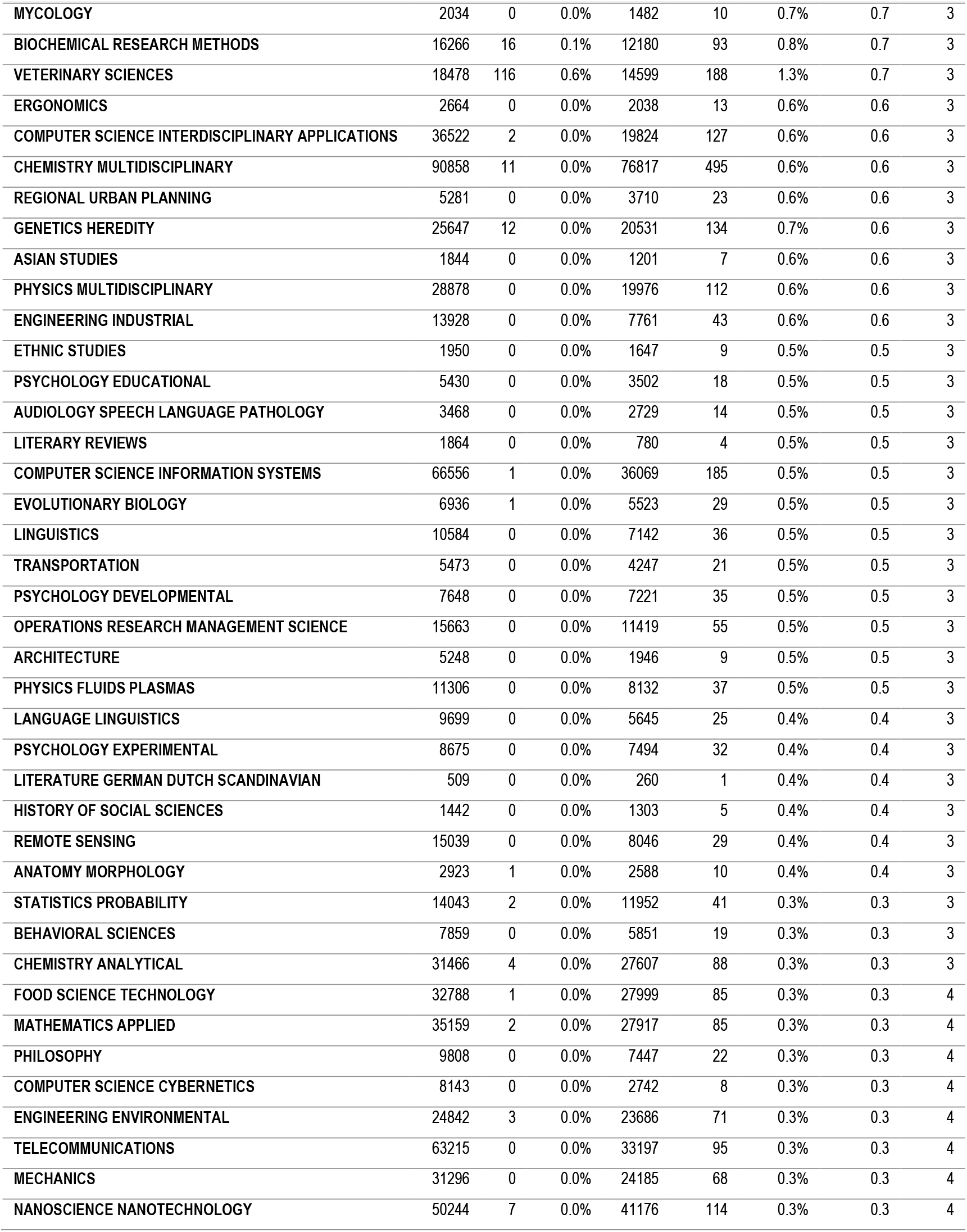
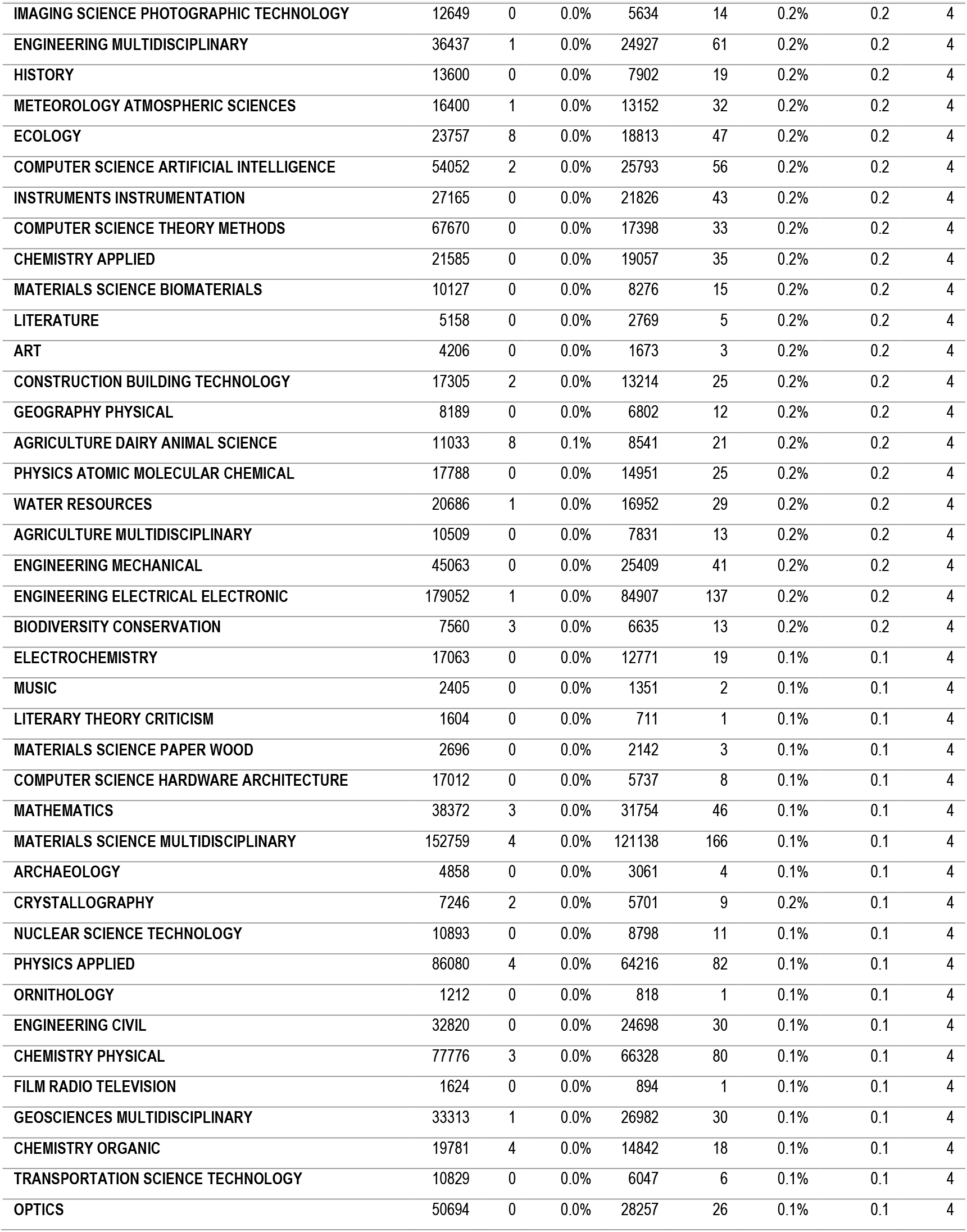
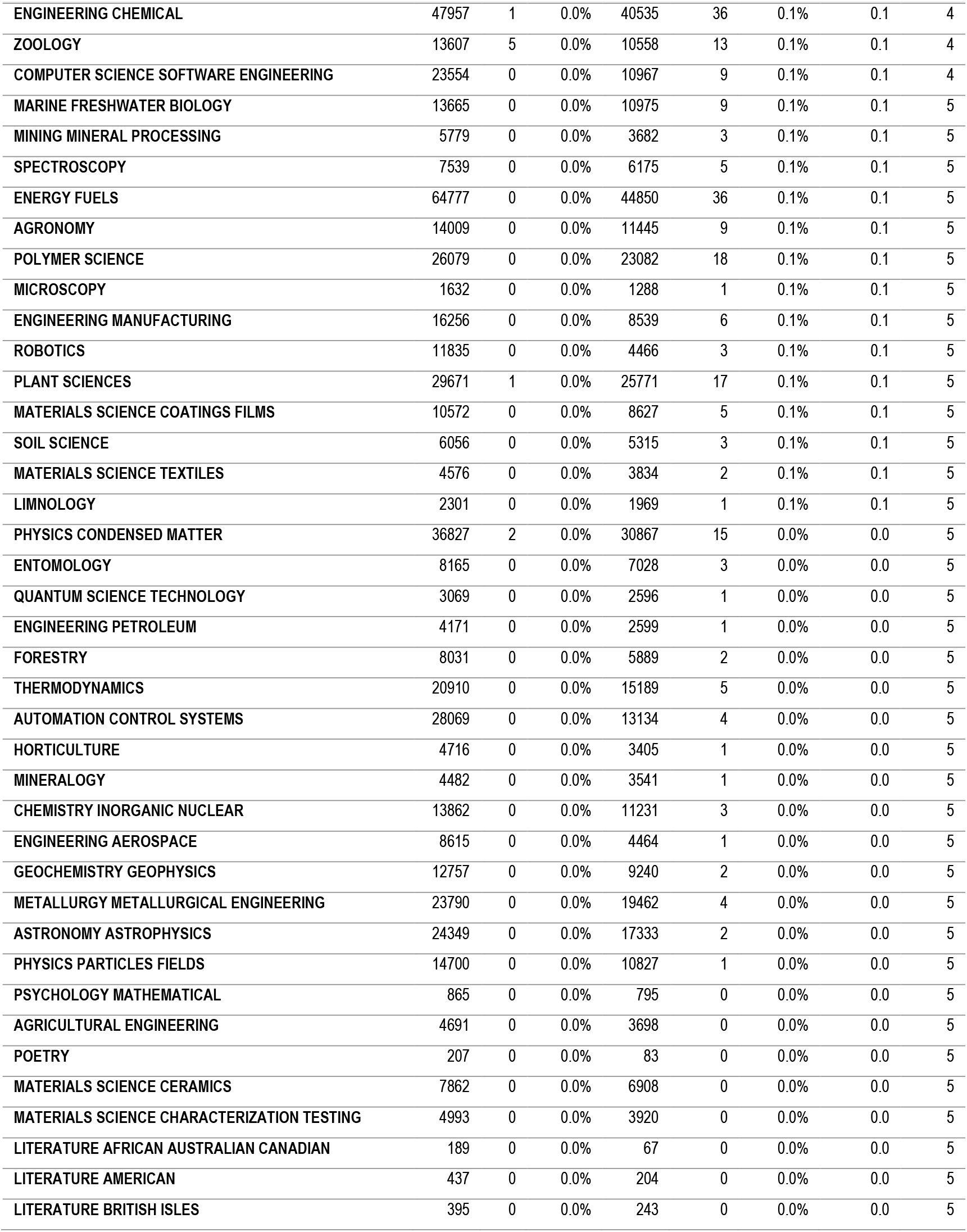
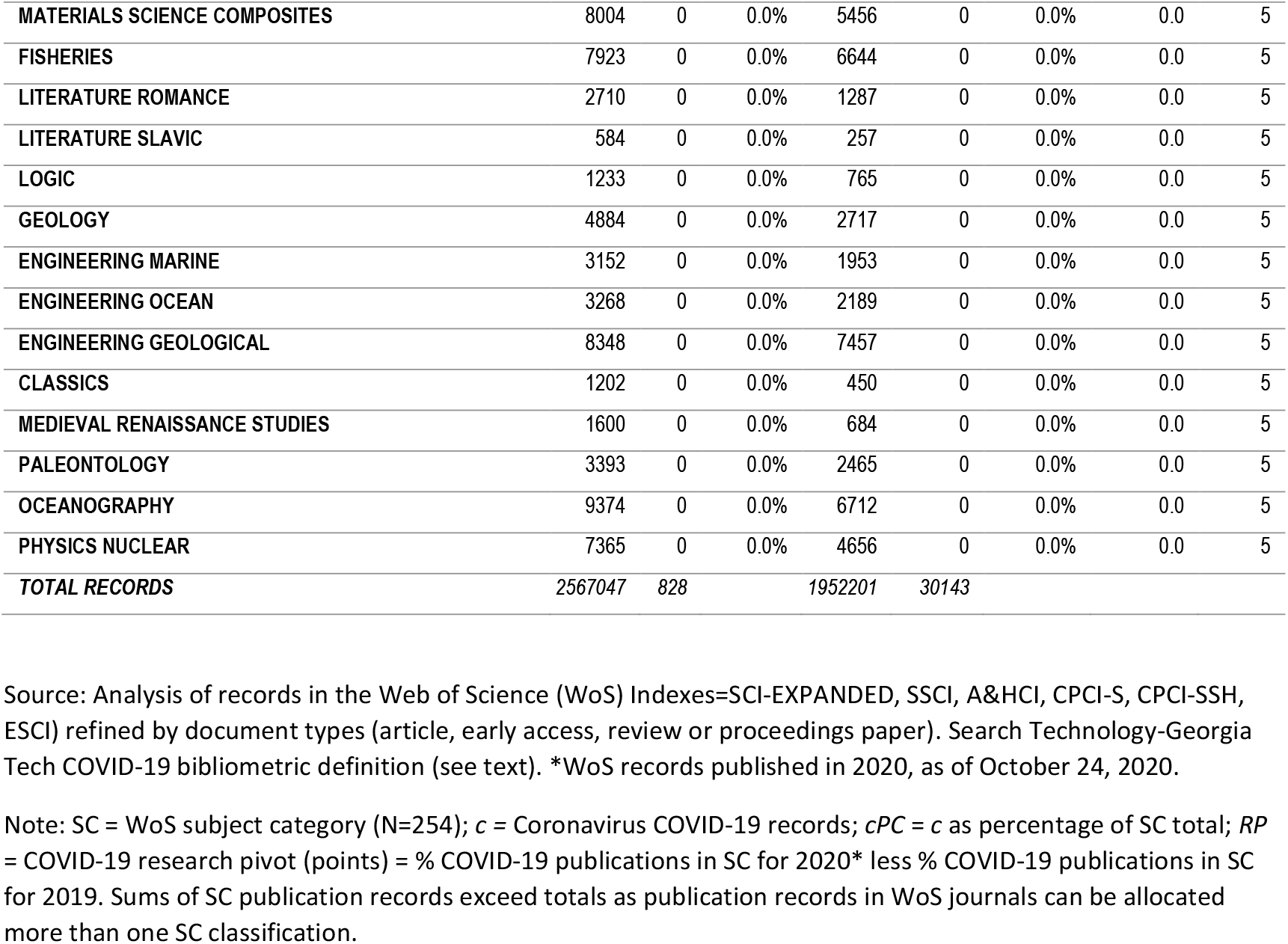

| Data Table for COVID-19 Research Pivot Calculations | WoS Publications<br>2019 |  |  | WoS Publications<br>2020* |  |  | Research<br>pivot | Quintile |
| --- | --- | --- | --- | --- | --- | --- | --- | --- |
| SUBJECT CATEGORY (SC) | Total | c | cPC | Total | c | cPC | RP |  |
| VIROLOGY | 6601 | 201 | 3.0% | 5703 | 1073 | 18.8% | 15.8 | 1 |
| MEDICAL ETHICS | 936 | 0 | 0.0% | 888 | 133 | 15.0% | 15.0 | 1 |
| INFECTIOUS DISEASES | 18110 | 132 | 0.7% | 15310 | 1559 | 10.2% | 9.5 | 1 |
| MEDICINE GENERAL INTERNAL | 49975 | 17 | 0.0% | 40534 | 2842 | 7.0% | 7.0 | 1 |
| OTORHINOLARYNGOLOGY | 7592 | 0 | 0.0% | 7036 | 453 | 6.4% | 6.4 | 1 |
| PUBLIC ADMINISTRATION | 3891 | 1 | 0.0% | 3819 | 240 | 6.3% | 6.3 | 1 |
| CRITICAL CARE MEDICINE | 5819 | 2 | 0.0% | 4878 | 274 | 5.6% | 5.6 | 1 |
| PUBLIC ENVIRONMENTAL OCCUPATIONAL HEALTH | 44853 | 80 | 0.2% | 39714 | 2171 | 5.5% | 5.3 | 1 |
| EMERGENCY MEDICINE | 4839 | 2 | 0.0% | 4165 | 221 | 5.3% | 5.3 | 1 |
| HEALTH CARE SCIENCES SERVICES | 17454 | 2 | 0.0% | 15560 | 803 | 5.2% | 5.1 | 1 |
| ANESTHESIOLOGY | 6213 | 0 | 0.0% | 5339 | 269 | 5.0% | 5.0 | 1 |
| TRANSPLANTATION | 4986 | 3 | 0.1% | 4465 | 225 | 5.0% | 5.0 | 1 |
| EDUCATION SCIENTIFIC DISCIPLINES | 10114 | 0 | 0.0% | 6407 | 312 | 4.9% | 4.9 | 1 |
| ETHICS | 3431 | 0 | 0.0% | 2974 | 141 | 4.7% | 4.7 | 1 |
| DERMATOLOGY | 9908 | 0 | 0.0% | 10322 | 473 | 4.6% | 4.6 | 1 |
| GERIATRICS GERONTOLOGY | 8633 | 0 | 0.0% | 8051 | 362 | 4.5% | 4.5 | 1 |
| IMMUNOLOGY | 27613 | 77 | 0.3% | 23464 | 1112 | 4.7% | 4.5 | 1 |
| PSYCHOLOGY CLINICAL | 11327 | 0 | 0.0% | 10685 | 476 | 4.5% | 4.5 | 1 |
| ALLERGY | 3230 | 2 | 0.1% | 2859 | 129 | 4.5% | 4.5 | 1 |
| RESPIRATORY SYSTEM | 11517 | 13 | 0.1% | 10112 | 458 | 4.5% | 4.4 | 1 |
| TROPICAL MEDICINE | 4475 | 7 | 0.2% | 3434 | 157 | 4.6% | 4.4 | 1 |
| HEMATOLOGY | 11848 | 5 | 0.0% | 11254 | 501 | 4.5% | 4.4 | 1 |
| SOCIAL ISSUES | 3174 | 0 | 0.0% | 2700 | 116 | 4.3% | 4.3 | 1 |
| GERONTOLOGY | 4848 | 0 | 0.0% | 4354 | 186 | 4.3% | 4.3 | 1 |
| PSYCHIATRY | 25184 | 1 | 0.0% | 23241 | 984 | 4.2% | 4.2 | 1 |
| PRIMARY HEALTH CARE | 1885 | 0 | 0.0% | 1577 | 66 | 4.2% | 4.2 | 1 |
| HEALTH POLICY SERVICES | 9055 | 4 | 0.0% | 8491 | 357 | 4.2% | 4.2 | 1 |
| PERIPHERAL VASCULAR DISEASE | 10671 | 0 | 0.0% | 9530 | 387 | 4.1% | 4.1 | 1 |
| SOCIAL SCIENCES BIOMEDICAL | 4284 | 0 | 0.0% | 4070 | 160 | 3.9% | 3.9 | 1 |
| CARDIAC CARDIOVASCULAR SYSTEMS | 24285 | 1 | 0.0% | 22664 | 888 | 3.9% | 3.9 | 1 |
| MEDICAL INFORMATICS | 7624 | 1 | 0.0% | 5395 | 208 | 3.9% | 3.8 | 1 |
| FOLKLORE | 277 | 0 | 0.0% | 183 | 7 | 3.8% | 3.8 | 1 |
| LAW | 11662 | 0 | 0.0% | 7288 | 277 | 3.8% | 3.8 | 1 |
| MEDICAL LABORATORY TECHNOLOGY | 3774 | 3 | 0.1% | 3116 | 119 | 3.8% | 3.7 | 1 |
| PEDIATRICS | 21278 | 11 | 0.1% | 20081 | 761 | 3.8% | 3.7 | 1 |
| SOCIAL WORK | 4894 | 0 | 0.0% | 4646 | 150 | 3.2% | 3.2 | 1 |
| MICROBIOLOGY | 26636 | 95 | 0.4% | 22645 | 805 | 3.6% | 3.2 | 1 |
| RADIOLOGY NUCLEAR MEDICINE MEDICAL IMAGING | 28828 | 1 | 0.0% | 23277 | 729 | 3.1% | 3.1 | 1 |
| WOMEN'S STUDIES | 3015 | 0 | 0.0% | 2532 | 78 | 3.1% | 3.1 | 1 |
| GEOGRAPHY | 8710 | 0 | 0.0% | 7040 | 210 | 3.0% | 3.0 | 1 |

| Data Table for COVID-19 Research Pivot Calculations |  |  |  | WoS Publications 2019 |  |  | WoS Publications 2020* |  |  | Research pivot | Quintile |
| --- | --- | --- | --- | --- | --- | --- | --- | --- | --- | --- | --- |
| SUBJECT CATEGORY (SC) |  |  |  | Total | c | cPC | Total | c | cPC | RP |  |
| COMMUNICATION |  |  |  | 9045 | 3 | 0.0% | 7089 | 210 | 3.0% | 2.9 | 1 |
| DENTISTRY ORAL SURGERY MEDICINE |  |  |  | 13330 | 0 | 0.0% | 11308 | 327 | 2.9% | 2.9 | 1 |
| MEDICINE RESEARCH EXPERIMENTAL |  |  |  | 36295 | 24 | 0.1% | 31110 | 917 | 2.9% | 2.9 | 1 |
| OBSTETRICS GYNECOLOGY |  |  |  | 16098 | 0 | 0.0% | 14578 | 420 | 2.9% | 2.9 | 1 |
| HOSPITALITY LEISURE SPORT TOURISM |  |  |  | 7611 | 0 | 0.0% | 7073 | 202 | 2.9% | 2.9 | 1 |
| GASTROENTEROLOGY HEPATOLOGY |  |  |  | 14830 | 1 | 0.0% | 13666 | 384 | 2.8% | 2.8 | 1 |
| UROLOGY NEPHROLOGY |  |  |  | 13074 | 0 | 0.0% | 11810 | 328 | 2.8% | 2.8 | 1 |
| RHEUMATOLOGY |  |  |  | 5906 | 1 | 0.0% | 5231 | 146 | 2.8% | 2.8 | 1 |
| MATHEMATICAL COMPUTATIONAL BIOLOGY |  |  |  | 10710 | 11 | 0.1% | 7301 | 210 | 2.9% | 2.8 | 1 |
| SURGERY |  |  |  | 46029 | 1 | 0.0% | 42330 | 1167 | 2.8% | 2.8 | 1 |
| INFORMATION SCIENCE LIBRARY SCIENCE |  |  |  | 8138 | 0 | 0.0% | 5856 | 161 | 2.7% | 2.7 | 1 |
| PARASITOLOGY |  |  |  | 6139 | 10 | 0.2% | 5364 | 151 | 2.8% | 2.7 | 2 |
| CLINICAL NEUROLOGY |  |  |  | 35934 | 0 | 0.0% | 31697 | 836 | 2.6% | 2.6 | 2 |
| AGRICULTURAL ECONOMICS POLICY |  |  |  | 2087 | 0 | 0.0% | 1385 | 36 | 2.6% | 2.6 | 2 |
| PSYCHOLOGY PSYCHOANALYSIS |  |  |  | 828 | 0 | 0.0% | 618 | 16 | 2.6% | 2.6 | 2 |
| AREA STUDIES |  |  |  | 5275 | 0 | 0.0% | 4208 | 108 | 2.6% | 2.6 | 2 |
| OPHTHALMOLOGY |  |  |  | 11695 | 1 | 0.0% | 10565 | 270 | 2.6% | 2.5 | 2 |
| BIOPHYSICS |  |  |  | 13335 | 11 | 0.1% | 11694 | 302 | 2.6% | 2.5 | 2 |
| PHARMACOLOGY PHARMACY |  |  |  | 54358 | 20 | 0.0% | 46974 | 1150 | 2.4% | 2.4 | 2 |
| SOCIOLOGY |  |  |  | 9674 | 0 | 0.0% | 8186 | 194 | 2.4% | 2.4 | 2 |
| NURSING |  |  |  | 13545 | 1 | 0.0% | 11814 | 267 | 2.3% | 2.3 | 2 |
| ANTHROPOLOGY |  |  |  | 5059 | 0 | 0.0% | 3836 | 86 | 2.2% | 2.2 | 2 |
| INTEGRATIVE COMPLEMENTARY MEDICINE |  |  |  | 4680 | 0 | 0.0% | 4021 | 89 | 2.2% | 2.2 | 2 |
| ENDOCRINOLOGY METABOLISM |  |  |  | 21829 | 0 | 0.0% | 19452 | 427 | 2.2% | 2.2 | 2 |
| PSYCHOLOGY SOCIAL |  |  |  | 5182 | 1 | 0.0% | 4553 | 96 | 2.1% | 2.1 | 2 |
| MEDICINE LEGAL |  |  |  | 2610 | 0 | 0.0% | 2074 | 43 | 2.1% | 2.1 | 2 |
| POLITICAL SCIENCE |  |  |  | 12328 | 1 | 0.0% | 10550 | 216 | 2.0% | 2.0 | 2 |
| INTERNATIONAL RELATIONS |  |  |  | 6677 | 0 | 0.0% | 5211 | 106 | 2.0% | 2.0 | 2 |
| CULTURAL STUDIES |  |  |  | 2412 | 0 | 0.0% | 1927 | 38 | 2.0% | 2.0 | 2 |
| SUBSTANCE ABUSE |  |  |  | 5302 | 0 | 0.0% | 4566 | 88 | 1.9% | 1.9 | 2 |
| CELL TISSUE ENGINEERING |  |  |  | 3459 | 0 | 0.0% | 2759 | 53 | 1.9% | 1.9 | 2 |
| SOCIAL SCIENCES INTERDISCIPLINARY |  |  |  | 15437 | 1 | 0.0% | 11171 | 212 | 1.9% | 1.9 | 2 |
| CELL BIOLOGY |  |  |  | 35833 | 19 | 0.1% | 30911 | 580 | 1.9% | 1.8 | 2 |
| NEUROIMAGING |  |  |  | 4020 | 0 | 0.0% | 2808 | 51 | 1.8% | 1.8 | 2 |
| REPRODUCTIVE BIOLOGY |  |  |  | 4672 | 0 | 0.0% | 3842 | 69 | 1.8% | 1.8 | 2 |
| ANDROLOGY |  |  |  | 818 | 0 | 0.0% | 1007 | 18 | 1.8% | 1.8 | 2 |
| PSYCHOLOGY MULTIDISCIPLINARY |  |  |  | 14271 | 0 | 0.0% | 12331 | 218 | 1.8% | 1.8 | 2 |
| ORTHOPEDICS |  |  |  | 16752 | 0 | 0.0% | 15059 | 253 | 1.7% | 1.7 | 2 |
| BIOLOGY |  |  |  | 18050 | 15 | 0.1% | 13688 | 241 | 1.8% | 1.7 | 2 |
| PSYCHOLOGY BIOLOGICAL |  |  |  | 1981 | 0 | 0.0% | 1637 | 27 | 1.6% | 1.6 | 2 |
| SUBJECT CATEGORY (SC) |  |  |  | Total | c | cPC | Total | c | cPC | RP |  |
| ONCOLOGY |  |  |  | 53955 | 5 | 0.0% | 48131 | 779 | 1.6% | 1.6 | 2 |
| URBAN STUDIES |  |  |  | 5788 | 0 | 0.0% | 4246 | 68 | 1.6% | 1.6 | 2 |
| MANAGEMENT |  |  |  | 25339 | 1 | 0.0% | 18229 | 291 | 1.6% | 1.6 | 2 |
| DANCE |  |  |  | 345 | 0 | 0.0% | 190 | 3 | 1.6% | 1.6 | 2 |
| RELIGION |  |  |  | 8768 | 0 | 0.0% | 5454 | 83 | 1.5% | 1.5 | 2 |
| PATHOLOGY |  |  |  | 9145 | 5 | 0.1% | 8159 | 124 | 1.5% | 1.5 | 2 |
| DEVELOPMENT STUDIES |  |  |  | 4451 | 0 | 0.0% | 3030 | 44 | 1.5% | 1.5 | 2 |
| ENVIRONMENTAL STUDIES |  |  |  | 20934 | 0 | 0.0% | 17470 | 246 | 1.4% | 1.4 | 2 |
| ECONOMICS |  |  |  | 35537 | 1 | 0.0% | 25325 | 350 | 1.4% | 1.4 | 2 |
| BIOCHEMISTRY MOLECULAR BIOLOGY |  |  |  | 69885 | 53 | 0.1% | 60269 | 876 | 1.5% | 1.4 | 2 |
| NEUROSCIENCES |  |  |  | 44989 | 1 | 0.0% | 37582 | 507 | 1.3% | 1.3 | 2 |
| HISTORY PHILOSOPHY OF SCIENCE |  |  |  | 3407 | 0 | 0.0% | 3197 | 43 | 1.3% | 1.3 | 2 |
| MULTIDISCIPLINARY SCIENCES |  |  |  | 75344 | 48 | 0.1% | 57678 | 795 | 1.4% | 1.3 | 2 |
| PSYCHOLOGY |  |  |  | 8471 | 0 | 0.0% | 6786 | 88 | 1.3% | 1.3 | 2 |
| BUSINESS |  |  |  | 18197 | 0 | 0.0% | 13820 | 179 | 1.3% | 1.3 | 2 |
| PSYCHOLOGY APPLIED |  |  |  | 5951 | 0 | 0.0% | 5520 | 71 | 1.3% | 1.3 | 2 |
| BUSINESS FINANCE |  |  |  | 10446 | 0 | 0.0% | 8385 | 106 | 1.3% | 1.3 | 2 |
| NUTRITION DIETETICS |  |  |  | 17404 | 2 | 0.0% | 14634 | 186 | 1.3% | 1.3 | 2 |
| PHYSIOLOGY |  |  |  | 13911 | 2 | 0.0% | 9969 | 126 | 1.3% | 1.2 | 2 |
| REHABILITATION |  |  |  | 12005 | 0 | 0.0% | 10231 | 125 | 1.2% | 1.2 | 2 |
| EDUCATION EDUCATIONAL RESEARCH |  |  |  | 44277 | 1 | 0.0% | 29614 | 361 | 1.2% | 1.2 | 2 |
| ENVIRONMENTAL SCIENCES |  |  |  | 89252 | 14 | 0.0% | 80916 | 946 | 1.2% | 1.2 | 2 |
| CHEMISTRY MEDICINAL |  |  |  | 19162 | 17 | 0.1% | 14005 | 167 | 1.2% | 1.1 | 3 |
| SOCIAL SCIENCES MATHEMATICAL METHODS |  |  |  | 3262 | 1 | 0.0% | 2856 | 31 | 1.1% | 1.1 | 3 |
| SPORT SCIENCES |  |  |  | 13238 | 1 | 0.0% | 10636 | 112 | 1.1% | 1.0 | 3 |
| ACOUSTICS |  |  |  | 8572 | 0 | 0.0% | 5239 | 54 | 1.0% | 1.0 | 3 |
| CRIMINOLOGY PENOLOGY |  |  |  | 5031 | 0 | 0.0% | 4875 | 47 | 1.0% | 1.0 | 3 |
| BIOTECHNOLOGY APPLIED MICROBIOLOGY |  |  |  | 31936 | 33 | 0.1% | 26572 | 259 | 1.0% | 0.9 | 3 |
| PHYSICS MATHEMATICAL |  |  |  | 11151 | 0 | 0.0% | 9372 | 78 | 0.8% | 0.8 | 3 |
| GREEN SUSTAINABLE SCIENCE TECHNOLOGY |  |  |  | 28992 | 0 | 0.0% | 22815 | 189 | 0.8% | 0.8 | 3 |
| THEATER |  |  |  | 1167 | 0 | 0.0% | 607 | 5 | 0.8% | 0.8 | 3 |
| MATHEMATICS INTERDISCIPLINARY APPLICATIONS |  |  |  | 14524 | 2 | 0.0% | 13276 | 105 | 0.8% | 0.8 | 3 |
| ENGINEERING BIOMEDICAL |  |  |  | 22982 | 1 | 0.0% | 13867 | 108 | 0.8% | 0.8 | 3 |
| EDUCATION SPECIAL |  |  |  | 2278 | 0 | 0.0% | 2230 | 17 | 0.8% | 0.8 | 3 |
| FAMILY STUDIES |  |  |  | 4580 | 0 | 0.0% | 4862 | 37 | 0.8% | 0.8 | 3 |
| HUMANITIES MULTIDISCIPLINARY |  |  |  | 11664 | 0 | 0.0% | 6647 | 50 | 0.8% | 0.8 | 3 |
| TOXICOLOGY |  |  |  | 13244 | 0 | 0.0% | 10786 | 81 | 0.8% | 0.8 | 3 |
| DEMOGRAPHY |  |  |  | 1991 | 0 | 0.0% | 1631 | 12 | 0.7% | 0.7 | 3 |
| DEVELOPMENTAL BIOLOGY |  |  |  | 3944 | 2 | 0.1% | 3636 | 28 | 0.8% | 0.7 | 3 |
| INDUSTRIAL RELATIONS LABOR |  |  |  | 1928 | 0 | 0.0% | 1594 | 11 | 0.7% | 0.7 | 3 |
| SUBJECT CATEGORY (SC) | Total | c | cPC | Total | c | cPC | RP |  |
| MYCOLOGY | 2034 | 0 | 0.0% | 1482 | 10 | 0.7% | 0.7 | 3 |
| BIOCHEMICAL RESEARCH METHODS | 16266 | 16 | 0.1% | 12180 | 93 | 0.8% | 0.7 | 3 |
| VETERINARY SCIENCES | 18478 | 116 | 0.6% | 14599 | 188 | 1.3% | 0.7 | 3 |
| ERGONOMICS | 2664 | 0 | 0.0% | 2038 | 13 | 0.6% | 0.6 | 3 |
| COMPUTER SCIENCE INTERDISCIPLINARY APPLICATIONS | 36522 | 2 | 0.0% | 19824 | 127 | 0.6% | 0.6 | 3 |
| CHEMISTRY MULTIDISCIPLINARY | 90858 | 11 | 0.0% | 76817 | 495 | 0.6% | 0.6 | 3 |
| REGIONAL URBAN PLANNING | 5281 | 0 | 0.0% | 3710 | 23 | 0.6% | 0.6 | 3 |
| GENETICS HEREDITY | 25647 | 12 | 0.0% | 20531 | 134 | 0.7% | 0.6 | 3 |
| ASIAN STUDIES | 1844 | 0 | 0.0% | 1201 | 7 | 0.6% | 0.6 | 3 |
| PHYSICS MULTIDISCIPLINARY | 28878 | 0 | 0.0% | 19976 | 112 | 0.6% | 0.6 | 3 |
| ENGINEERING INDUSTRIAL | 13928 | 0 | 0.0% | 7761 | 43 | 0.6% | 0.6 | 3 |
| ETHNIC STUDIES | 1950 | 0 | 0.0% | 1647 | 9 | 0.5% | 0.5 | 3 |
| PSYCHOLOGY EDUCATIONAL | 5430 | 0 | 0.0% | 3502 | 18 | 0.5% | 0.5 | 3 |
| AUDIOLOGY SPEECH LANGUAGE PATHOLOGY | 3468 | 0 | 0.0% | 2729 | 14 | 0.5% | 0.5 | 3 |
| LITERARY REVIEWS | 1864 | 0 | 0.0% | 780 | 4 | 0.5% | 0.5 | 3 |
| COMPUTER SCIENCE INFORMATION SYSTEMS | 66556 | 1 | 0.0% | 36069 | 185 | 0.5% | 0.5 | 3 |
| EVOLUTIONARY BIOLOGY | 6936 | 1 | 0.0% | 5523 | 29 | 0.5% | 0.5 | 3 |
| LINGUISTICS | 10584 | 0 | 0.0% | 7142 | 36 | 0.5% | 0.5 | 3 |
| TRANSPORTATION | 5473 | 0 | 0.0% | 4247 | 21 | 0.5% | 0.5 | 3 |
| PSYCHOLOGY DEVELOPMENTAL | 7648 | 0 | 0.0% | 7221 | 35 | 0.5% | 0.5 | 3 |
| OPERATIONS RESEARCH MANAGEMENT SCIENCE | 15663 | 0 | 0.0% | 11419 | 55 | 0.5% | 0.5 | 3 |
| ARCHITECTURE | 5248 | 0 | 0.0% | 1946 | 9 | 0.5% | 0.5 | 3 |
| PHYSICS FLUIDS PLASMAS | 11306 | 0 | 0.0% | 8132 | 37 | 0.5% | 0.5 | 3 |
| LANGUAGE LINGUISTICS | 9699 | 0 | 0.0% | 5645 | 25 | 0.4% | 0.4 | 3 |
| PSYCHOLOGY EXPERIMENTAL | 8675 | 0 | 0.0% | 7494 | 32 | 0.4% | 0.4 | 3 |
| LITERATURE GERMAN DUTCH SCANDINAVIAN | 509 | 0 | 0.0% | 260 | 1 | 0.4% | 0.4 | 3 |
| HISTORY OF SOCIAL SCIENCES | 1442 | 0 | 0.0% | 1303 | 5 | 0.4% | 0.4 | 3 |
| REMOTE SENSING | 15039 | 0 | 0.0% | 8046 | 29 | 0.4% | 0.4 | 3 |
| ANATOMY MORPHOLOGY | 2923 | 1 | 0.0% | 2588 | 10 | 0.4% | 0.4 | 3 |
| STATISTICS PROBABILITY | 14043 | 2 | 0.0% | 11952 | 41 | 0.3% | 0.3 | 3 |
| BEHAVIORAL SCIENCES | 7859 | 0 | 0.0% | 5851 | 19 | 0.3% | 0.3 | 3 |
| CHEMISTRY ANALYTICAL | 31466 | 4 | 0.0% | 27607 | 88 | 0.3% | 0.3 | 3 |
| FOOD SCIENCE TECHNOLOGY | 32788 | 1 | 0.0% | 27999 | 85 | 0.3% | 0.3 | 4 |
| MATHEMATICS APPLIED | 35159 | 2 | 0.0% | 27917 | 85 | 0.3% | 0.3 | 4 |
| PHILOSOPHY | 9808 | 0 | 0.0% | 7447 | 22 | 0.3% | 0.3 | 4 |
| COMPUTER SCIENCE CYBERNETICS | 8143 | 0 | 0.0% | 2742 | 8 | 0.3% | 0.3 | 4 |
| ENGINEERING ENVIRONMENTAL | 24842 | 3 | 0.0% | 23686 | 71 | 0.3% | 0.3 | 4 |
| TELECOMMUNICATIONS | 63215 | 0 | 0.0% | 33197 | 95 | 0.3% | 0.3 | 4 |
| MECHANICS | 31296 | 0 | 0.0% | 24185 | 68 | 0.3% | 0.3 | 4 |
| NANOSCIENCE NANOTECHNOLOGY | 50244 | 7 | 0.0% | 41176 | 114 | 0.3% | 0.3 | 4 |
| SUBJECT CATEGORY (SC) | Total | c | cPC | Total | c | cPC | RP |  |
| IMAGING SCIENCE PHOTOGRAPHIC TECHNOLOGY | 12649 | 0 | 0.0% | 5634 | 14 | 0.2% | 0.2 | 4 |
| ENGINEERING MULTIDISCIPLINARY | 36437 | 1 | 0.0% | 24927 | 61 | 0.2% | 0.2 | 4 |
| HISTORY | 13600 | 0 | 0.0% | 7902 | 19 | 0.2% | 0.2 | 4 |
| METEOROLOGY ATMOSPHERIC SCIENCES | 16400 | 1 | 0.0% | 13152 | 32 | 0.2% | 0.2 | 4 |
| ECOLOGY | 23757 | 8 | 0.0% | 18813 | 47 | 0.2% | 0.2 | 4 |
| COMPUTER SCIENCE ARTIFICIAL INTELLIGENCE | 54052 | 2 | 0.0% | 25793 | 56 | 0.2% | 0.2 | 4 |
| INSTRUMENTS INSTRUMENTATION | 27165 | 0 | 0.0% | 21826 | 43 | 0.2% | 0.2 | 4 |
| COMPUTER SCIENCE THEORY METHODS | 67670 | 0 | 0.0% | 17398 | 33 | 0.2% | 0.2 | 4 |
| CHEMISTRY APPLIED | 21585 | 0 | 0.0% | 19057 | 35 | 0.2% | 0.2 | 4 |
| MATERIALS SCIENCE BIOMATERIALS | 10127 | 0 | 0.0% | 8276 | 15 | 0.2% | 0.2 | 4 |
| LITERATURE | 5158 | 0 | 0.0% | 2769 | 5 | 0.2% | 0.2 | 4 |
| ART | 4206 | 0 | 0.0% | 1673 | 3 | 0.2% | 0.2 | 4 |
| CONSTRUCTION BUILDING TECHNOLOGY | 17305 | 2 | 0.0% | 13214 | 25 | 0.2% | 0.2 | 4 |
| GEOGRAPHY PHYSICAL | 8189 | 0 | 0.0% | 6802 | 12 | 0.2% | 0.2 | 4 |
| AGRICULTURE DAIRY ANIMAL SCIENCE | 11033 | 8 | 0.1% | 8541 | 21 | 0.2% | 0.2 | 4 |
| PHYSICS ATOMIC MOLECULAR CHEMICAL | 17788 | 0 | 0.0% | 14951 | 25 | 0.2% | 0.2 | 4 |
| WATER RESOURCES | 20686 | 1 | 0.0% | 16952 | 29 | 0.2% | 0.2 | 4 |
| AGRICULTURE MULTIDISCIPLINARY | 10509 | 0 | 0.0% | 7831 | 13 | 0.2% | 0.2 | 4 |
| ENGINEERING MECHANICAL | 45063 | 0 | 0.0% | 25409 | 41 | 0.2% | 0.2 | 4 |
| ENGINEERING ELECTRICAL ELECTRONIC | 179052 | 1 | 0.0% | 84907 | 137 | 0.2% | 0.2 | 4 |
| BIODIVERSITY CONSERVATION | 7560 | 3 | 0.0% | 6635 | 13 | 0.2% | 0.2 | 4 |
| ELECTROCHEMISTRY | 17063 | 0 | 0.0% | 12771 | 19 | 0.1% | 0.1 | 4 |
| MUSIC | 2405 | 0 | 0.0% | 1351 | 2 | 0.1% | 0.1 | 4 |
| LITERARY THEORY CRITICISM | 1604 | 0 | 0.0% | 711 | 1 | 0.1% | 0.1 | 4 |
| MATERIALS SCIENCE PAPER WOOD | 2696 | 0 | 0.0% | 2142 | 3 | 0.1% | 0.1 | 4 |
| COMPUTER SCIENCE HARDWARE ARCHITECTURE | 17012 | 0 | 0.0% | 5737 | 8 | 0.1% | 0.1 | 4 |
| MATHEMATICS | 38372 | 3 | 0.0% | 31754 | 46 | 0.1% | 0.1 | 4 |
| MATERIALS SCIENCE MULTIDISCIPLINARY | 152759 | 4 | 0.0% | 121138 | 166 | 0.1% | 0.1 | 4 |
| ARCHAEOLOGY | 4858 | 0 | 0.0% | 3061 | 4 | 0.1% | 0.1 | 4 |
| CRYSTALLOGRAPHY | 7246 | 2 | 0.0% | 5701 | 9 | 0.2% | 0.1 | 4 |
| NUCLEAR SCIENCE TECHNOLOGY | 10893 | 0 | 0.0% | 8798 | 11 | 0.1% | 0.1 | 4 |
| PHYSICS APPLIED | 86080 | 4 | 0.0% | 64216 | 82 | 0.1% | 0.1 | 4 |
| ORNITHOLOGY | 1212 | 0 | 0.0% | 818 | 1 | 0.1% | 0.1 | 4 |
| ENGINEERING CIVIL | 32820 | 0 | 0.0% | 24698 | 30 | 0.1% | 0.1 | 4 |
| CHEMISTRY PHYSICAL | 77776 | 3 | 0.0% | 66328 | 80 | 0.1% | 0.1 | 4 |
| FILM RADIO TELEVISION | 1624 | 0 | 0.0% | 894 | 1 | 0.1% | 0.1 | 4 |
| GEOSCIENCES MULTIDISCIPLINARY | 33313 | 1 | 0.0% | 26982 | 30 | 0.1% | 0.1 | 4 |
| CHEMISTRY ORGANIC | 19781 | 4 | 0.0% | 14842 | 18 | 0.1% | 0.1 | 4 |
| TRANSPORTATION SCIENCE TECHNOLOGY | 10829 | 0 | 0.0% | 6047 | 6 | 0.1% | 0.1 | 4 |
| OPTICS | 50694 | 0 | 0.0% | 28257 | 26 | 0.1% | 0.1 | 4 |
| SUBJECT CATEGORY (SC) | Total | c | cPC | Total | c | cPC | RP |  |
| ENGINEERING CHEMICAL | 47957 | 1 | 0.0% | 40535 | 36 | 0.1% | 0.1 | 4 |
| ZOOLOGY | 13607 | 5 | 0.0% | 10558 | 13 | 0.1% | 0.1 | 4 |
| COMPUTER SCIENCE SOFTWARE ENGINEERING | 23554 | 0 | 0.0% | 10967 | 9 | 0.1% | 0.1 | 4 |
| MARINE FRESHWATER BIOLOGY | 13665 | 0 | 0.0% | 10975 | 9 | 0.1% | 0.1 | 5 |
| MINING MINERAL PROCESSING | 5779 | 0 | 0.0% | 3682 | 3 | 0.1% | 0.1 | 5 |
| SPECTROSCOPY | 7539 | 0 | 0.0% | 6175 | 5 | 0.1% | 0.1 | 5 |
| ENERGY FUELS | 64777 | 0 | 0.0% | 44850 | 36 | 0.1% | 0.1 | 5 |
| AGRONOMY | 14009 | 0 | 0.0% | 11445 | 9 | 0.1% | 0.1 | 5 |
| POLYMER SCIENCE | 26079 | 0 | 0.0% | 23082 | 18 | 0.1% | 0.1 | 5 |
| MICROSCOPY | 1632 | 0 | 0.0% | 1288 | 1 | 0.1% | 0.1 | 5 |
| ENGINEERING MANUFACTURING | 16256 | 0 | 0.0% | 8539 | 6 | 0.1% | 0.1 | 5 |
| ROBOTICS | 11835 | 0 | 0.0% | 4466 | 3 | 0.1% | 0.1 | 5 |
| PLANT SCIENCES | 29671 | 1 | 0.0% | 25771 | 17 | 0.1% | 0.1 | 5 |
| MATERIALS SCIENCE COATINGS FILMS | 10572 | 0 | 0.0% | 8627 | 5 | 0.1% | 0.1 | 5 |
| SOIL SCIENCE | 6056 | 0 | 0.0% | 5315 | 3 | 0.1% | 0.1 | 5 |
| MATERIALS SCIENCE TEXTILES | 4576 | 0 | 0.0% | 3834 | 2 | 0.1% | 0.1 | 5 |
| LIMNOLOGY | 2301 | 0 | 0.0% | 1969 | 1 | 0.1% | 0.1 | 5 |
| PHYSICS CONDENSED MATTER | 36827 | 2 | 0.0% | 30867 | 15 | 0.0% | 0.0 | 5 |
| ENTOMOLOGY | 8165 | 0 | 0.0% | 7028 | 3 | 0.0% | 0.0 | 5 |
| QUANTUM SCIENCE TECHNOLOGY | 3069 | 0 | 0.0% | 2596 | 1 | 0.0% | 0.0 | 5 |
| ENGINEERING PETROLEUM | 4171 | 0 | 0.0% | 2599 | 1 | 0.0% | 0.0 | 5 |
| FORESTRY | 8031 | 0 | 0.0% | 5889 | 2 | 0.0% | 0.0 | 5 |
| THERMODYNAMICS | 20910 | 0 | 0.0% | 15189 | 5 | 0.0% | 0.0 | 5 |
| AUTOMATION CONTROL SYSTEMS | 28069 | 0 | 0.0% | 13134 | 4 | 0.0% | 0.0 | 5 |
| HORTICULTURE | 4716 | 0 | 0.0% | 3405 | 1 | 0.0% | 0.0 | 5 |
| MINERALOGY | 4482 | 0 | 0.0% | 3541 | 1 | 0.0% | 0.0 | 5 |
| CHEMISTRY INORGANIC NUCLEAR | 13862 | 0 | 0.0% | 11231 | 3 | 0.0% | 0.0 | 5 |
| ENGINEERING AEROSPACE | 8615 | 0 | 0.0% | 4464 | 1 | 0.0% | 0.0 | 5 |
| GEOCHEMISTRY GEOPHYSICS | 12757 | 0 | 0.0% | 9240 | 2 | 0.0% | 0.0 | 5 |
| METALLURGY METALLURGICAL ENGINEERING | 23790 | 0 | 0.0% | 19462 | 4 | 0.0% | 0.0 | 5 |
| ASTRONOMY ASTROPHYSICS | 24349 | 0 | 0.0% | 17333 | 2 | 0.0% | 0.0 | 5 |
| PHYSICS PARTICLES FIELDS | 14700 | 0 | 0.0% | 10827 | 1 | 0.0% | 0.0 | 5 |
| PSYCHOLOGY MATHEMATICAL | 865 | 0 | 0.0% | 795 | 0 | 0.0% | 0.0 | 5 |
| AGRICULTURAL ENGINEERING | 4691 | 0 | 0.0% | 3698 | 0 | 0.0% | 0.0 | 5 |
| POETRY | 207 | 0 | 0.0% | 83 | 0 | 0.0% | 0.0 | 5 |
| MATERIALS SCIENCE CERAMICS | 7862 | 0 | 0.0% | 6908 | 0 | 0.0% | 0.0 | 5 |
| MATERIALS SCIENCE CHARACTERIZATION TESTING | 4993 | 0 | 0.0% | 3920 | 0 | 0.0% | 0.0 | 5 |
| LITERATURE AFRICAN AUSTRALIAN CANADIAN | 189 | 0 | 0.0% | 67 | 0 | 0.0% | 0.0 | 5 |
| LITERATURE AMERICAN | 437 | 0 | 0.0% | 204 | 0 | 0.0% | 0.0 | 5 |
| LITERATURE BRITISH ISLES | 395 | 0 | 0.0% | 243 | 0 | 0.0% | 0.0 | 5 |
| SUBJECT CATEGORY (SC) | Total | c | cPC | Total | c | cPC | RP |  |
| MATERIALS SCIENCE COMPOSITES | 8004 | 0 | 0.0% | 5456 | 0 | 0.0% | 0.0 | 5 |
| FISHERIES | 7923 | 0 | 0.0% | 6644 | 0 | 0.0% | 0.0 | 5 |
| LITERATURE ROMANCE | 2710 | 0 | 0.0% | 1287 | 0 | 0.0% | 0.0 | 5 |
| LITERATURE SLAVIC | 584 | 0 | 0.0% | 257 | 0 | 0.0% | 0.0 | 5 |
| LOGIC | 1233 | 0 | 0.0% | 765 | 0 | 0.0% | 0.0 | 5 |
| GEOLOGY | 4884 | 0 | 0.0% | 2717 | 0 | 0.0% | 0.0 | 5 |
| ENGINEERING MARINE | 3152 | 0 | 0.0% | 1953 | 0 | 0.0% | 0.0 | 5 |
| ENGINEERING OCEAN | 3268 | 0 | 0.0% | 2189 | 0 | 0.0% | 0.0 | 5 |
| ENGINEERING GEOLOGICAL | 8348 | 0 | 0.0% | 7457 | 0 | 0.0% | 0.0 | 5 |
| CLASSICS | 1202 | 0 | 0.0% | 450 | 0 | 0.0% | 0.0 | 5 |
| MEDIEVAL RENAISSANCE STUDIES | 1600 | 0 | 0.0% | 684 | 0 | 0.0% | 0.0 | 5 |
| PALEONTOLOGY | 3393 | 0 | 0.0% | 2465 | 0 | 0.0% | 0.0 | 5 |
| OCEANOGRAPHY | 9374 | 0 | 0.0% | 6712 | 0 | 0.0% | 0.0 | 5 |
| PHYSICS NUCLEAR | 7365 | 0 | 0.0% | 4656 | 0 | 0.0% | 0.0 | 5 |
| TOTAL RECORDS | 2567047 | 828 |  | 1952201 | 30143 |  |  |  |

Source: Analysis of records in the Web of Science (WoS) Indexes=SCI-EXPANDED, SSCI, A&HCI, CPCI-S, CPCI-SSH, ESCI) refined by document types (article, early access, review or proceedings paper). Search Technology-Georgia Tech COVID-19 bibliometric definition (see text). \*WoS records published in 2020, as of October 24, 2020.
Note: SC = WoS subject category (N=254); c = Coronavirus COVID-19 records; cPC = c as percentage of SC total; RP = COVID-19 research pivot (points) = % COVID-19 publications in SC for 2020\* less % COVID-19 publications in SC for 2019. Sums of SC publication records exceed totals as publication records in WoS journals can be allocated more than one SC classification.

## Appendix 2

This appendix provides an updated bibliometric analysis through to mid-April 2022. This updated was added on April 27, 2022.

The definition used for searching for research publications on COVID-19 topics remains the same as used in the original analysis. In addition to the Web of Science and PubMed, the appendix reports results from the Dimensions database of publications (https://www.dimensions.ai/). For discussion and calculation method of COVID-19 Research Pivot points, see text of main working paper.

### Contents of Appendix 2

Table 1 Overall Publication Trends, Web of Science, Dimensions, and PubMed, 2019-2022*

Table 2 COVID 19 publications reported in the Web of Science (WoS), 2019-2022*

Table 3 COVID-19 Publications by Top 30 Publishing Countries, Web of Science (WoS), 2019-2022*

Table 4 COVID 19 publications (including preprints) reported in Dimensions, 2019-2022*

### Sources and Notes

The tables in this appendix should be read in conjunction with these notes:

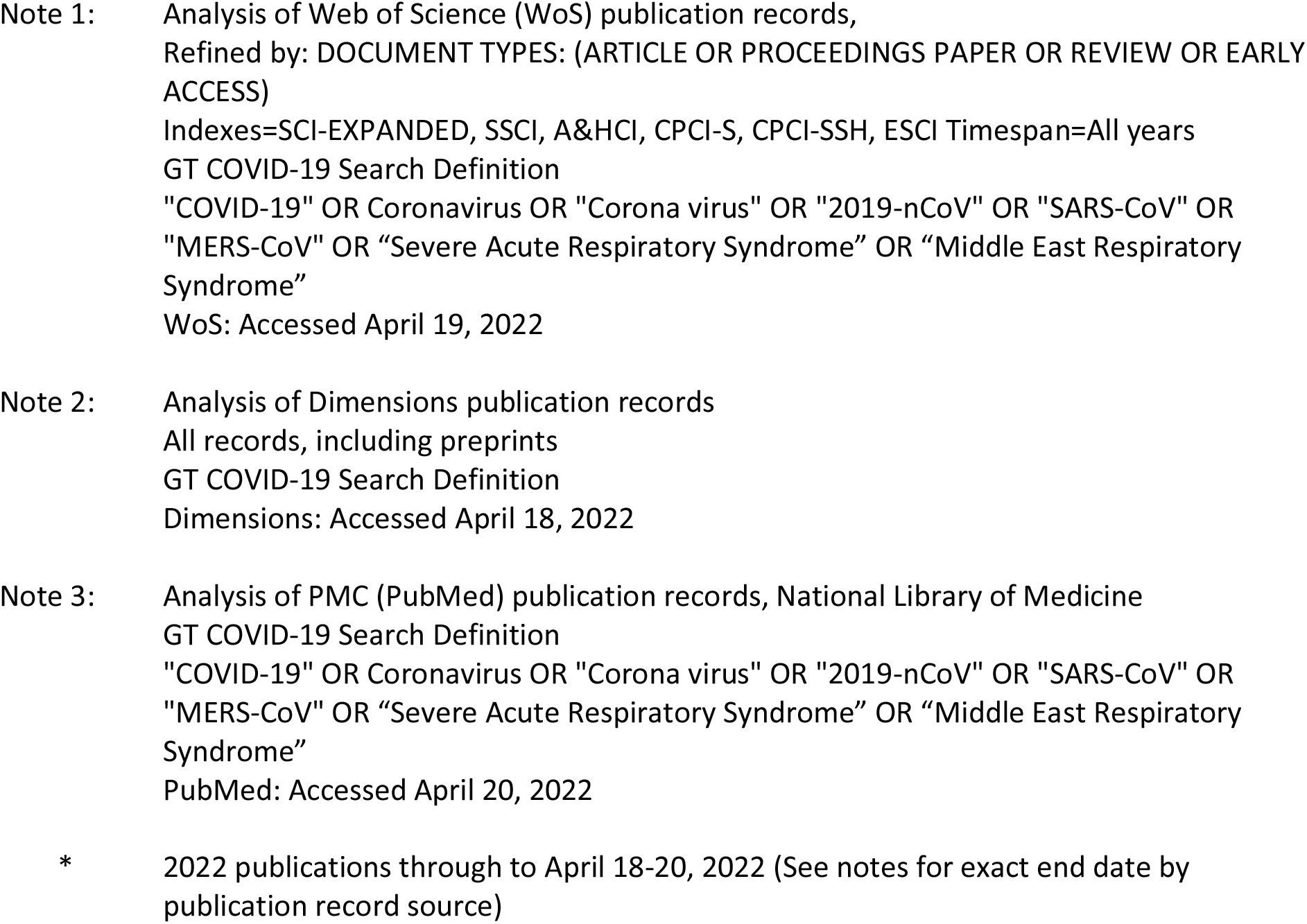

**Appendix 2. Table 1.**
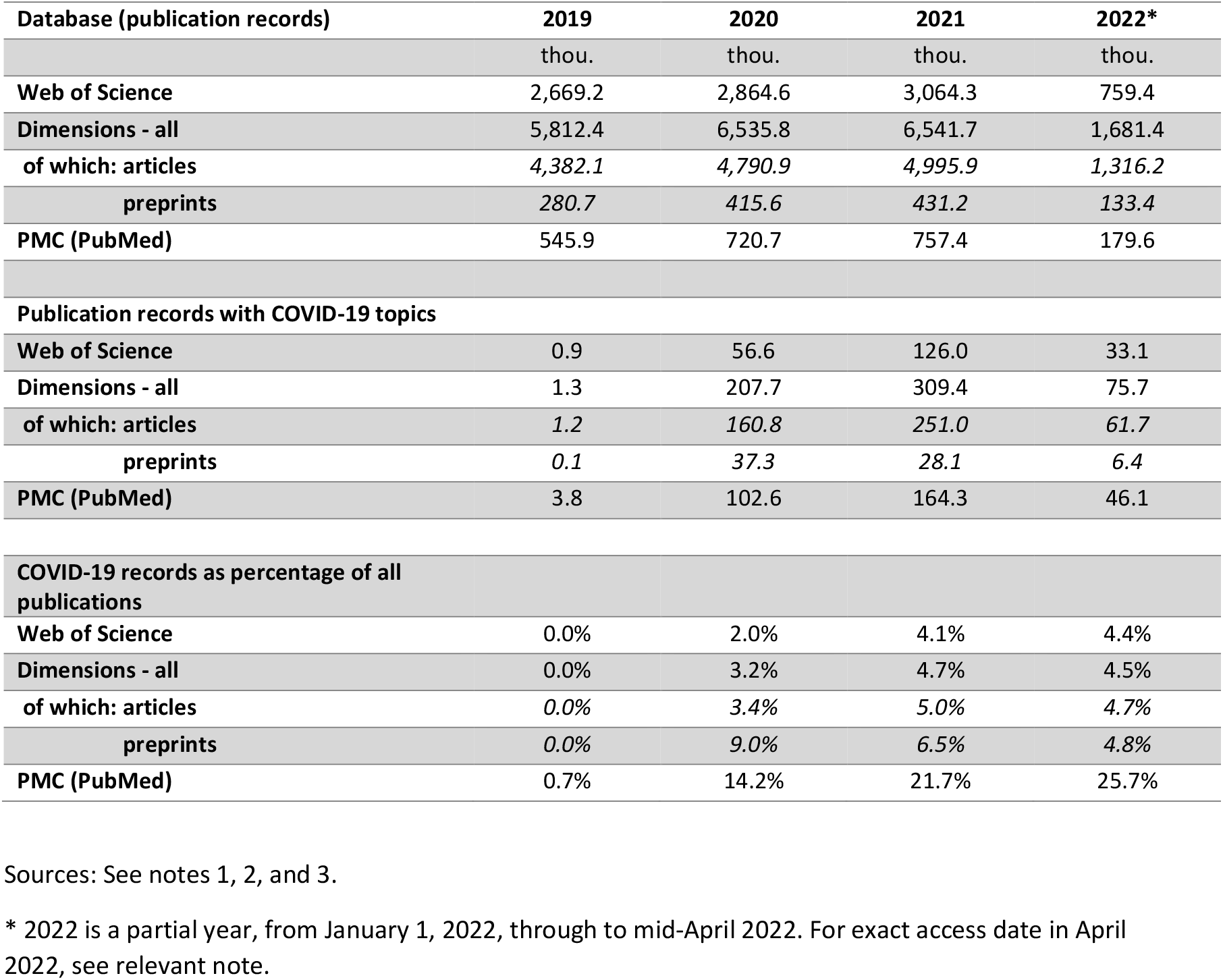
Overall Publication Trends, Web of Science, Dimensions, and PubMed, 2019-2022*.

| **Database (publication records)** | **2019** | **2020** | **2021** | **2022\*** |
| --- | --- | --- | --- | --- |
|  | thou. | thou. | thou. | thou. |
| **Web of Science** | 2,669.2 | 2,864.6 | 3,064.3 | 759.4 |
| **Dimensions - all** | 5,812.4 | 6,535.8 | 6,541.7 | 1,681.4 |
| **of which: articles** | 4,382.1 | 4,790.9 | 4,995.9 | 1,316.2 |
| **preprints** | 280.7 | 415.6 | 431.2 | 133.4 |
| **PMC (PubMed)** | 545.9 | 720.7 | 757.4 | 179.6 |
| **Publication records with COVID-19 topics** |  |  |  |  |
| **Web of Science** | 0.9 | 56.6 | 126.0 | 33.1 |
| **Dimensions - all** | 1.3 | 207.7 | 309.4 | 75.7 |
| **of which: articles** | 1.2 | 160.8 | 251.0 | 61.7 |
| **preprints** | 0.1 | 37.3 | 28.1 | 6.4 |
| **PMC (PubMed)** | 3.8 | 102.6 | 164.3 | 46.1 |
| **COVID-19 records as percentage of all publications** |  |  |  |  |
| **Web of Science** | 0.0% | 2.0% | 4.1% | 4.4% |
| **Dimensions - all** | 0.0% | 3.2% | 4.7% | 4.5% |
| **of which: articles** | 0.0% | 3.4% | 5.0% | 4.7% |
| **preprints** | 0.0% | 9.0% | 6.5% | 4.8% |
| **PMC (PubMed)** | 0.7% | 14.2% | 21.7% | 25.7% |

Sources: See notes 1, 2, and 3.
\* 2022 is a partial year, from January 1, 2022, through to mid-April 2022. For exact access date in April 2022, see relevant note.

**Appendix 2. Table 2.**
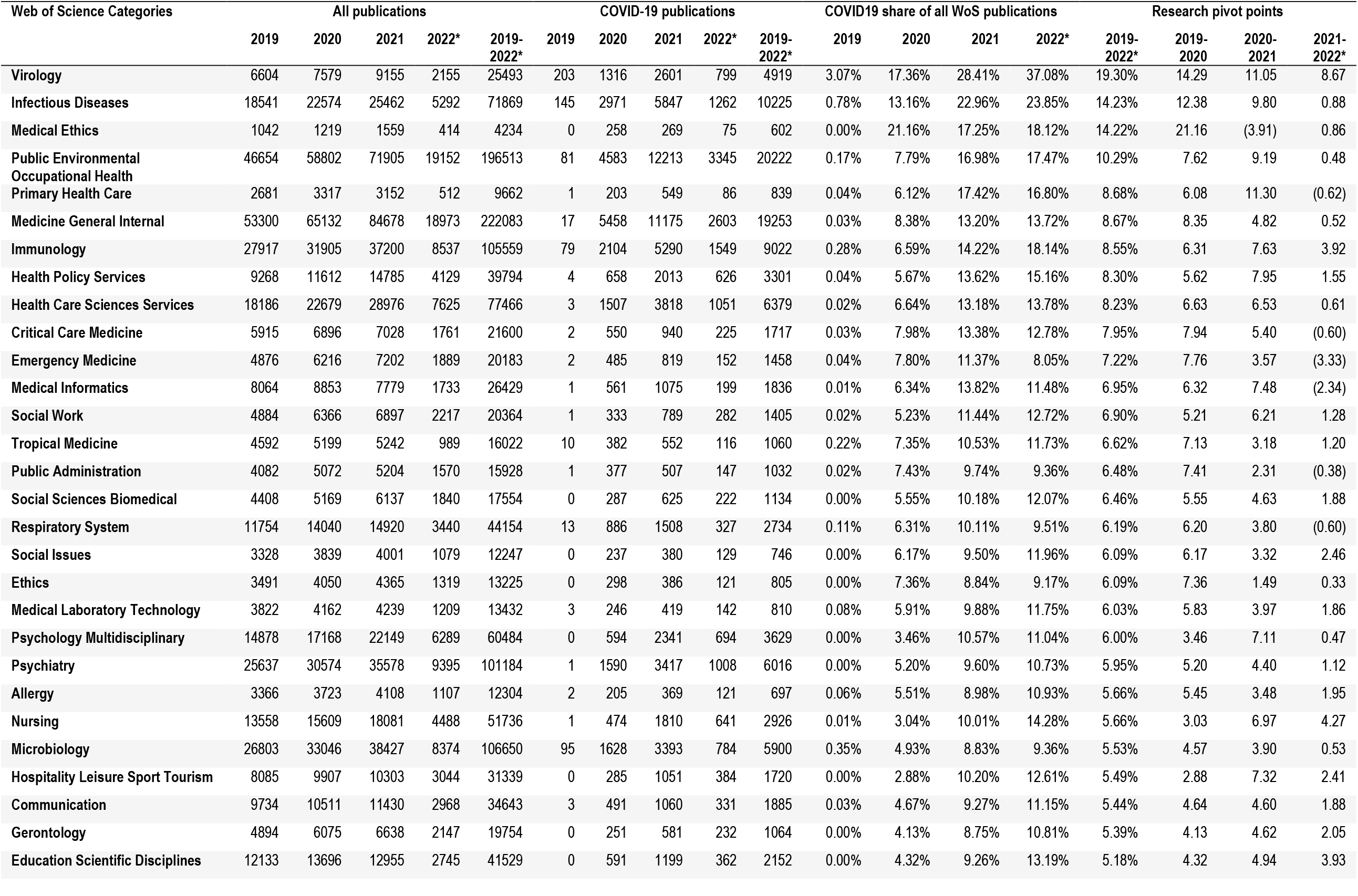
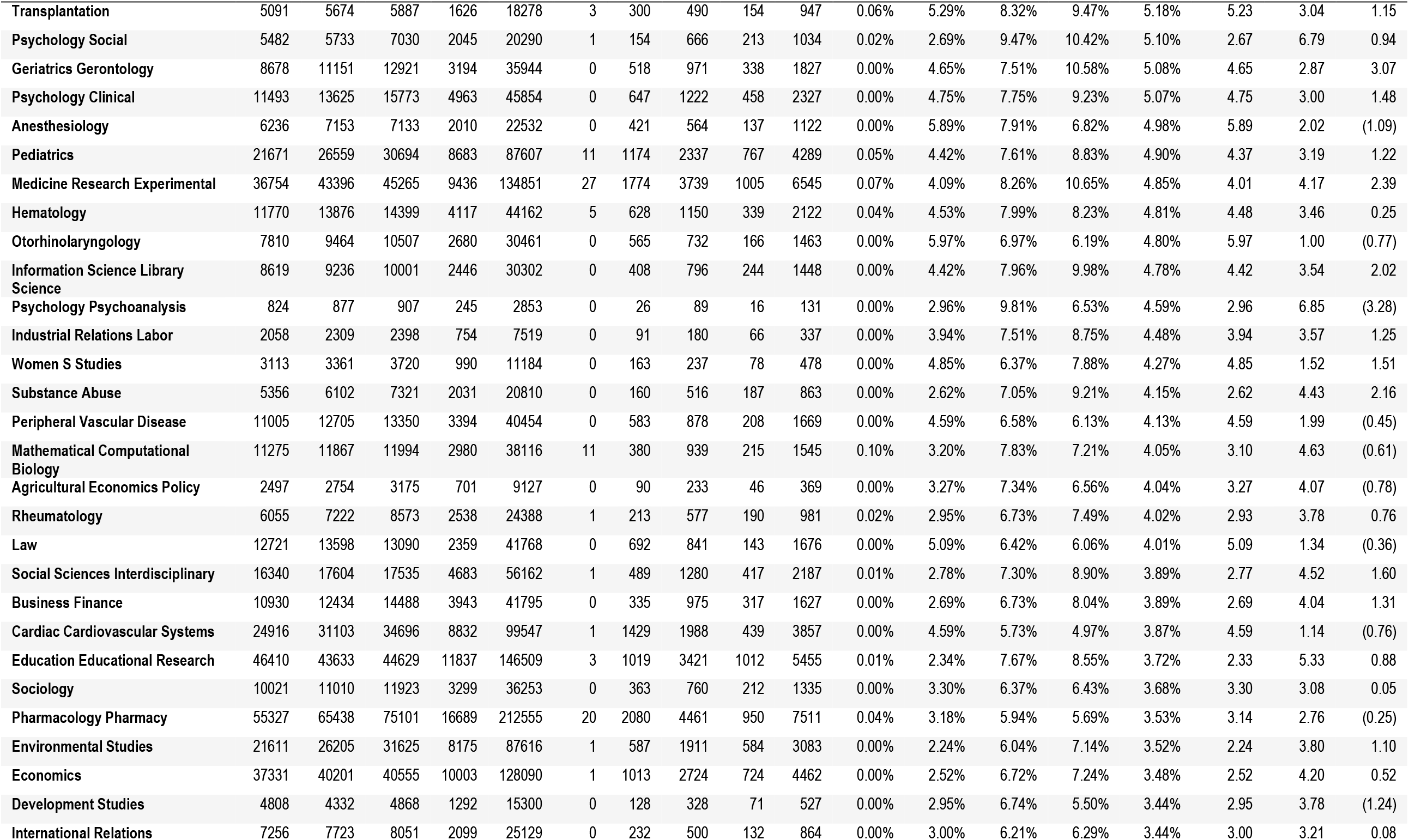
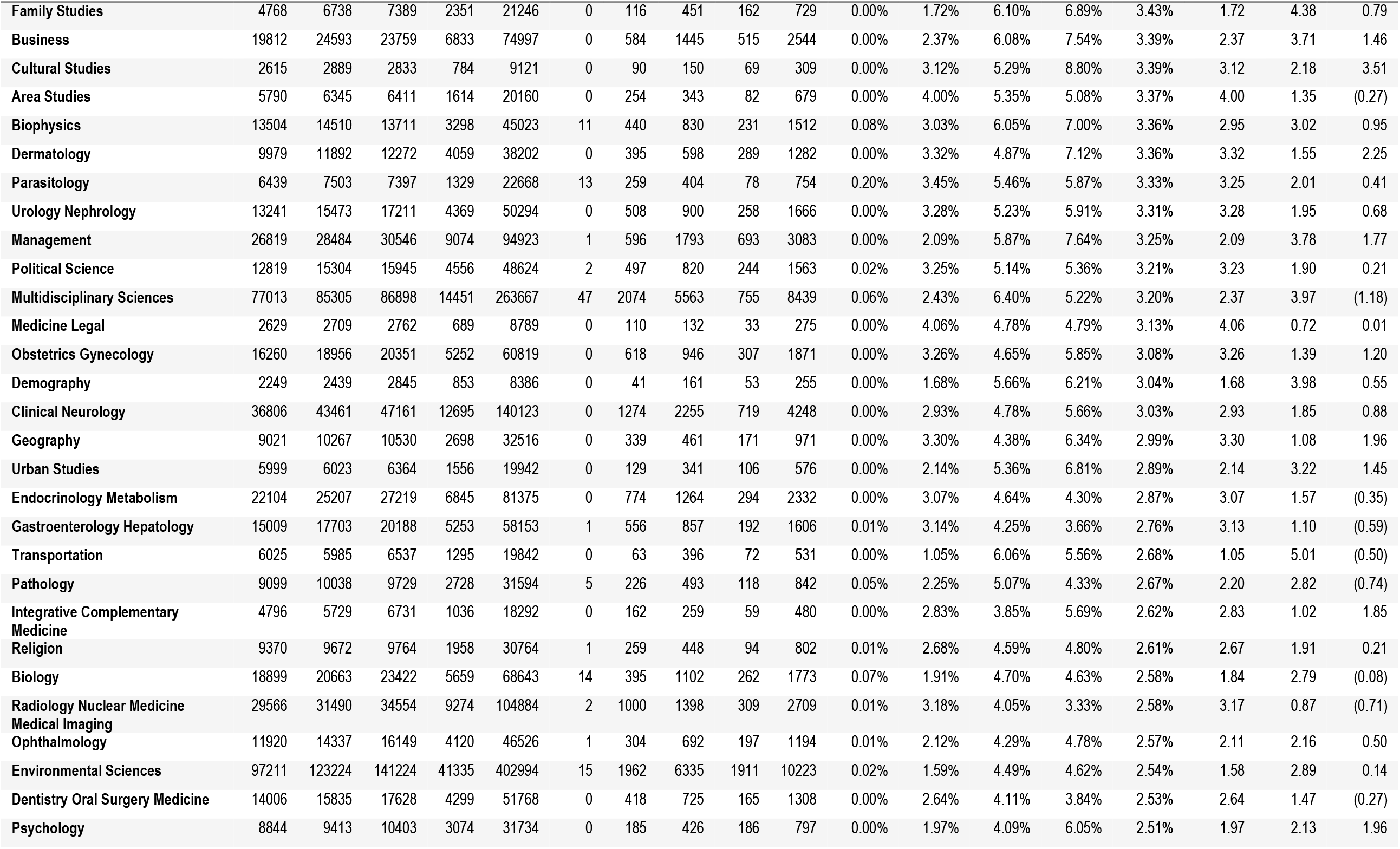
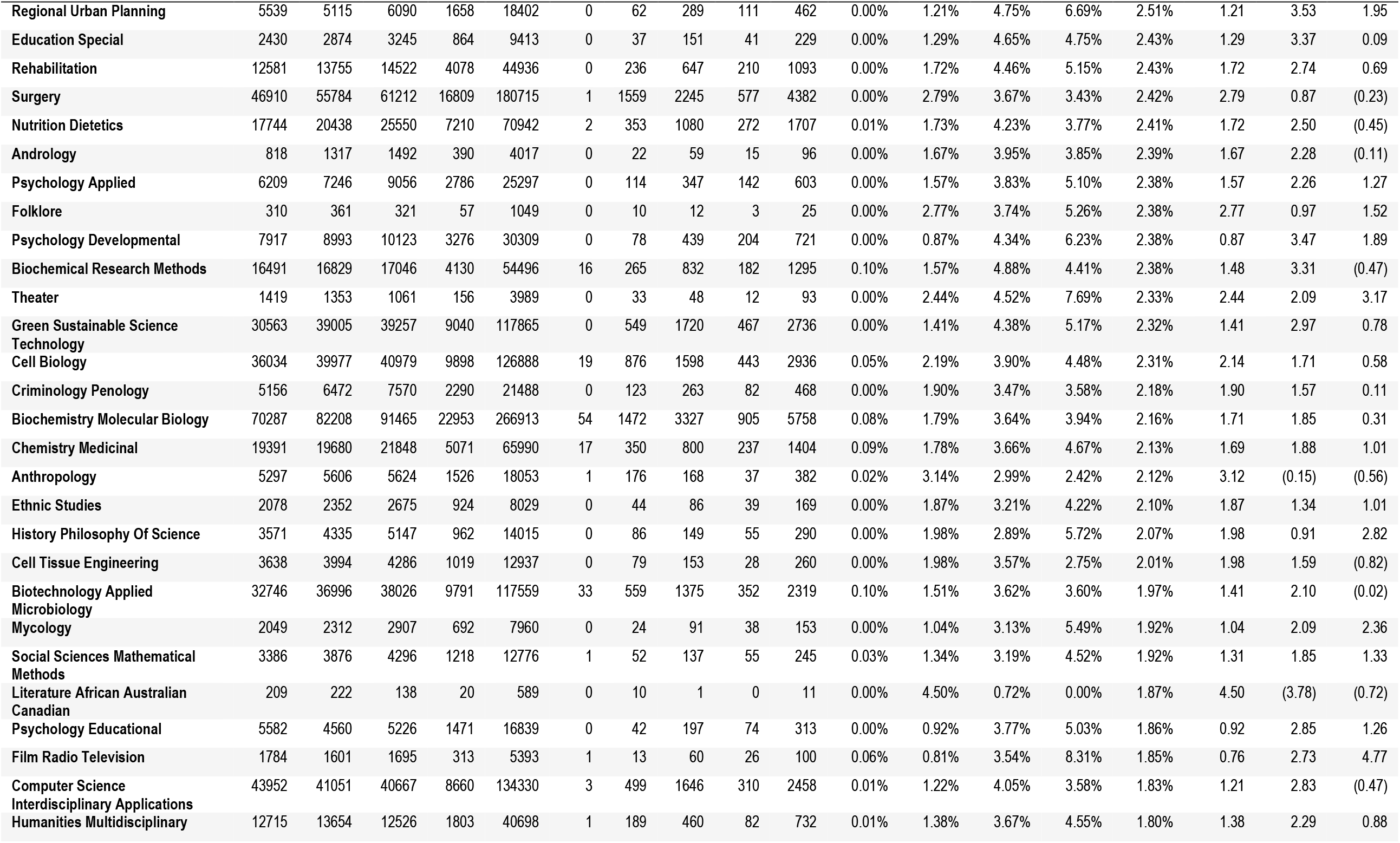
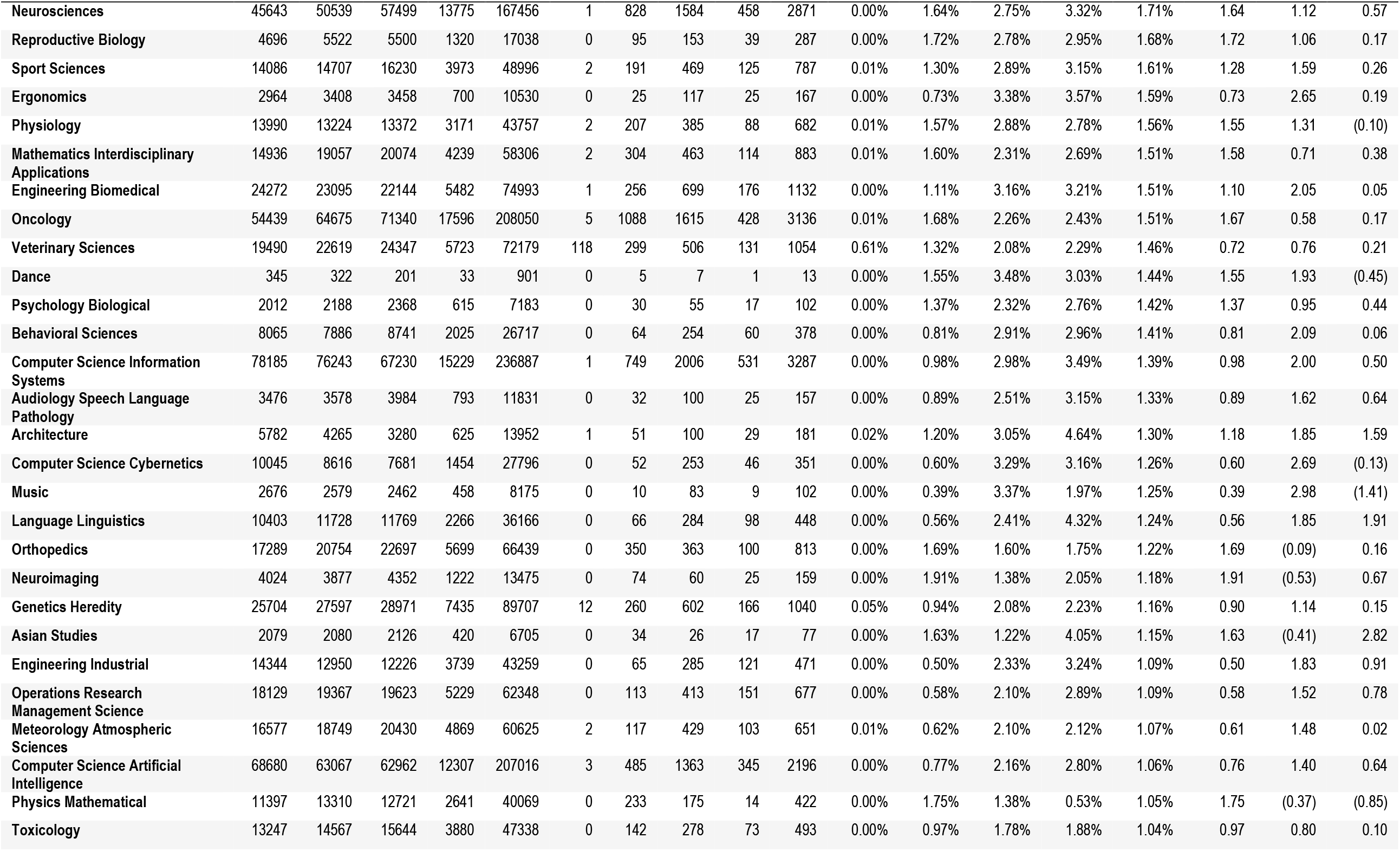
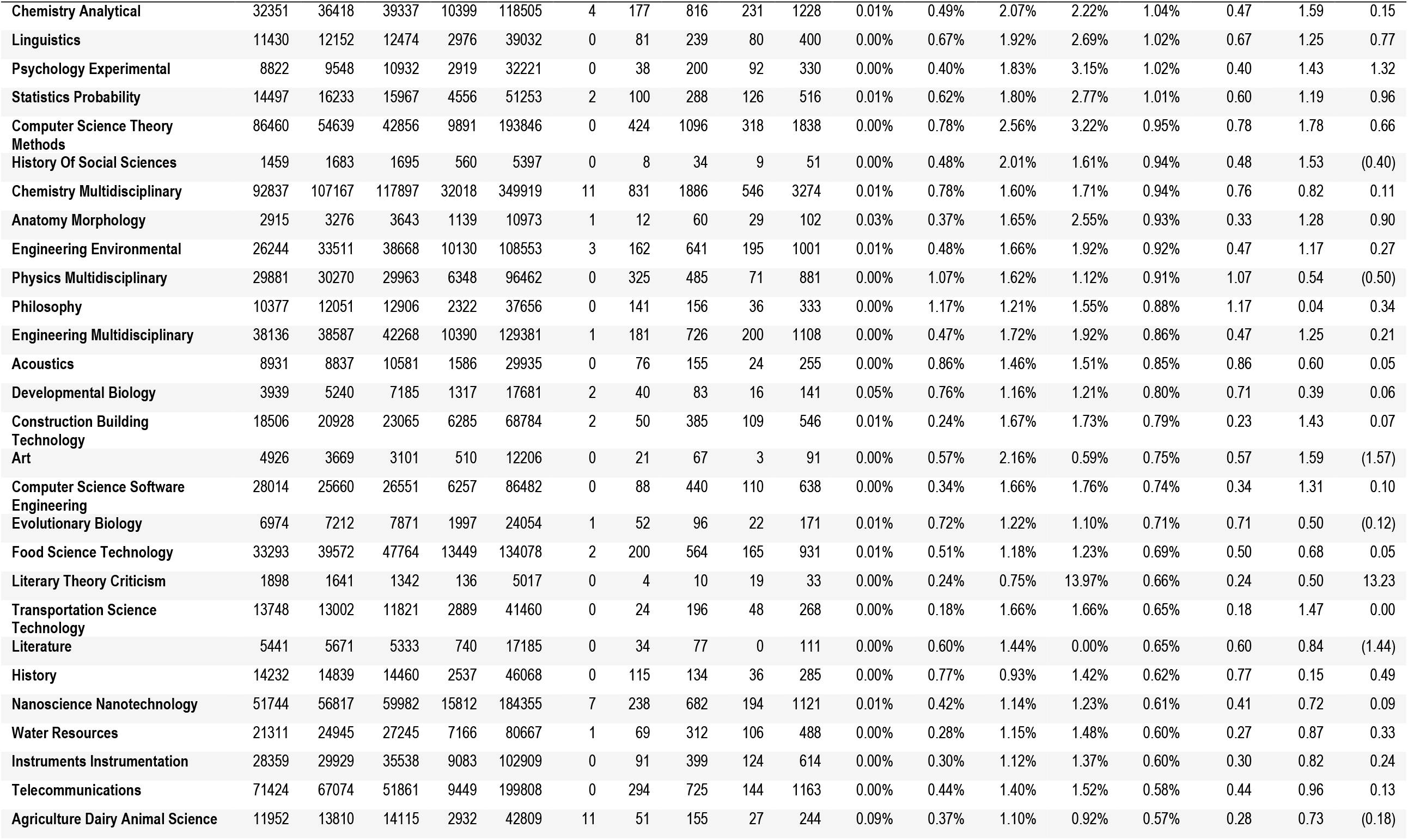
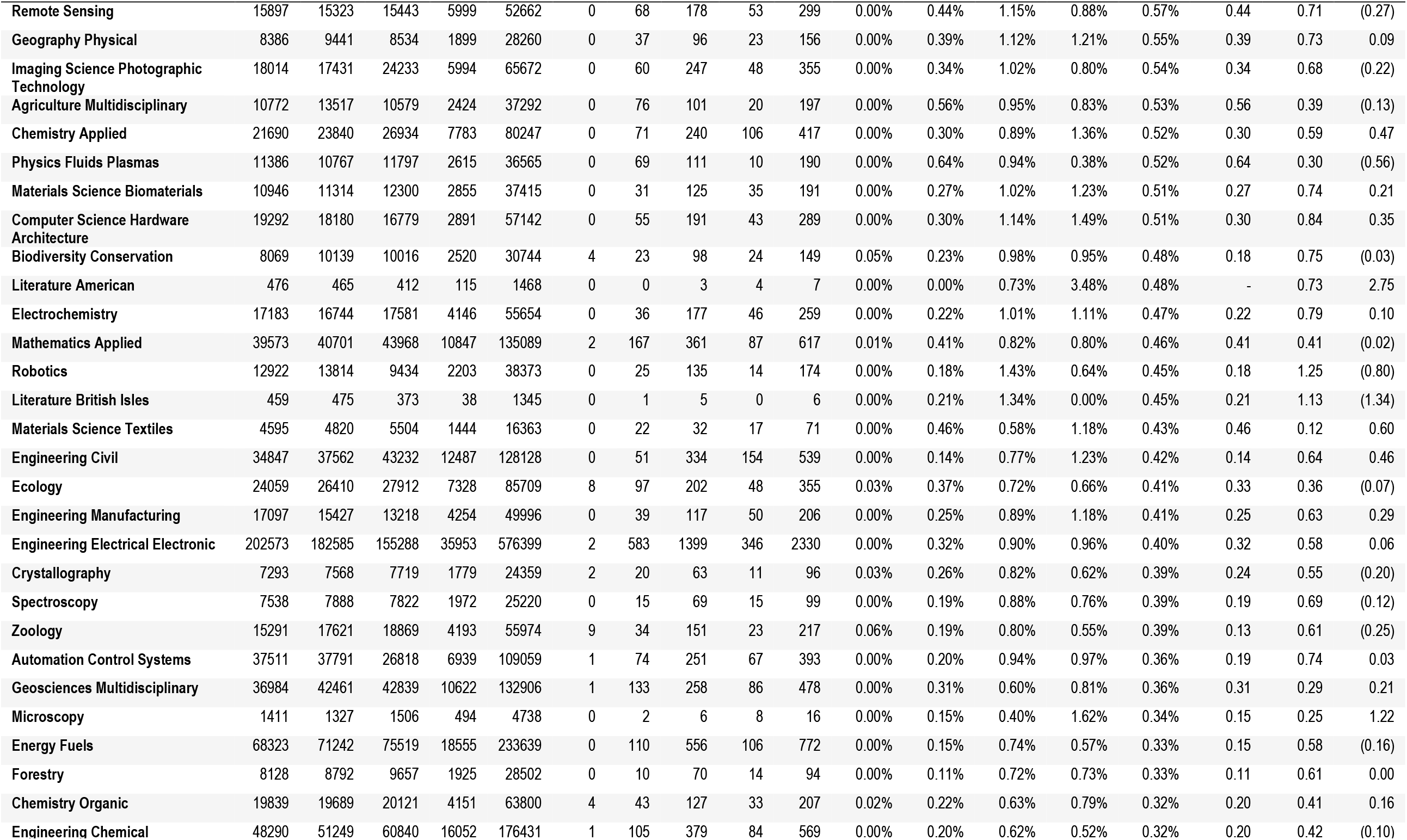
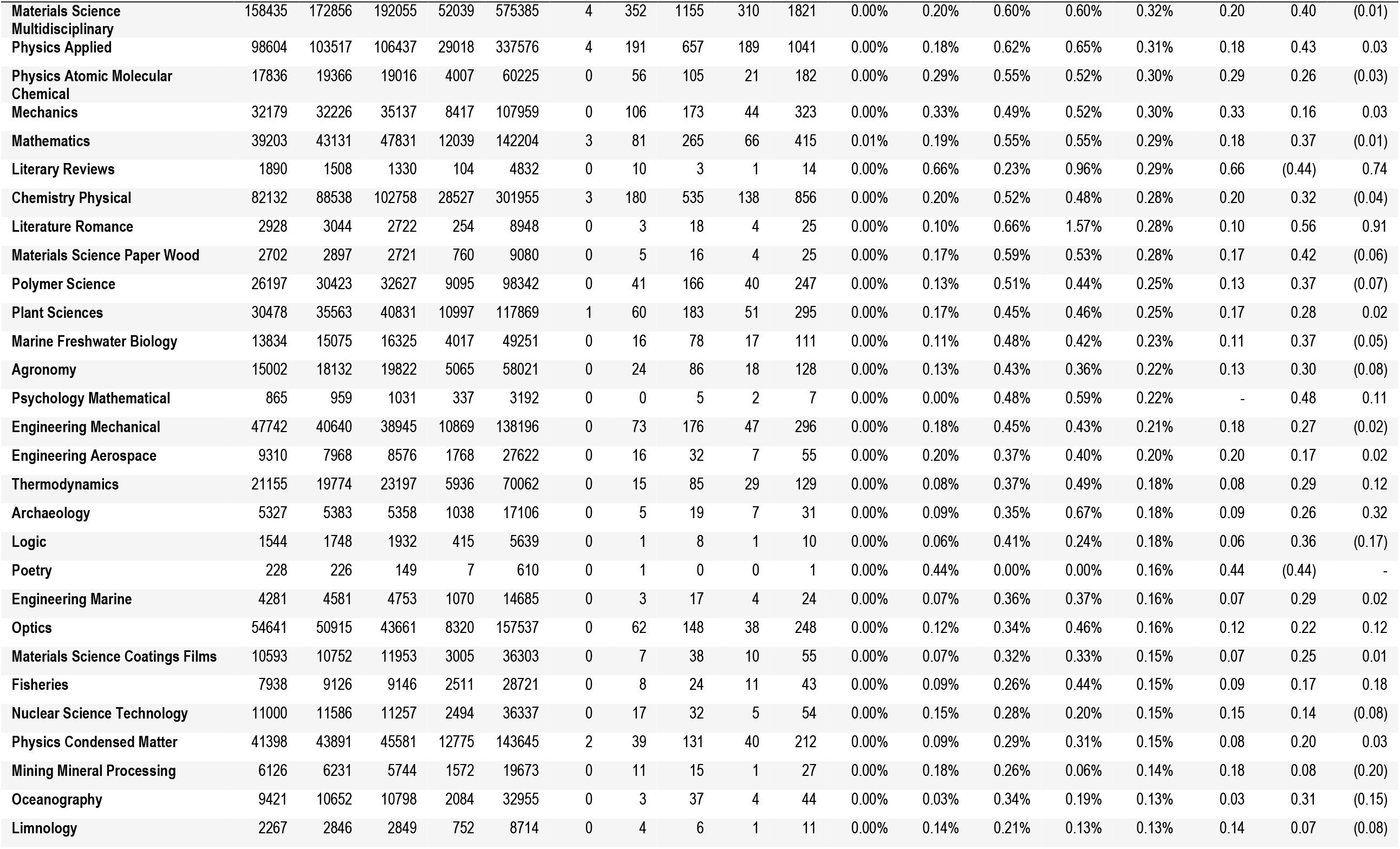
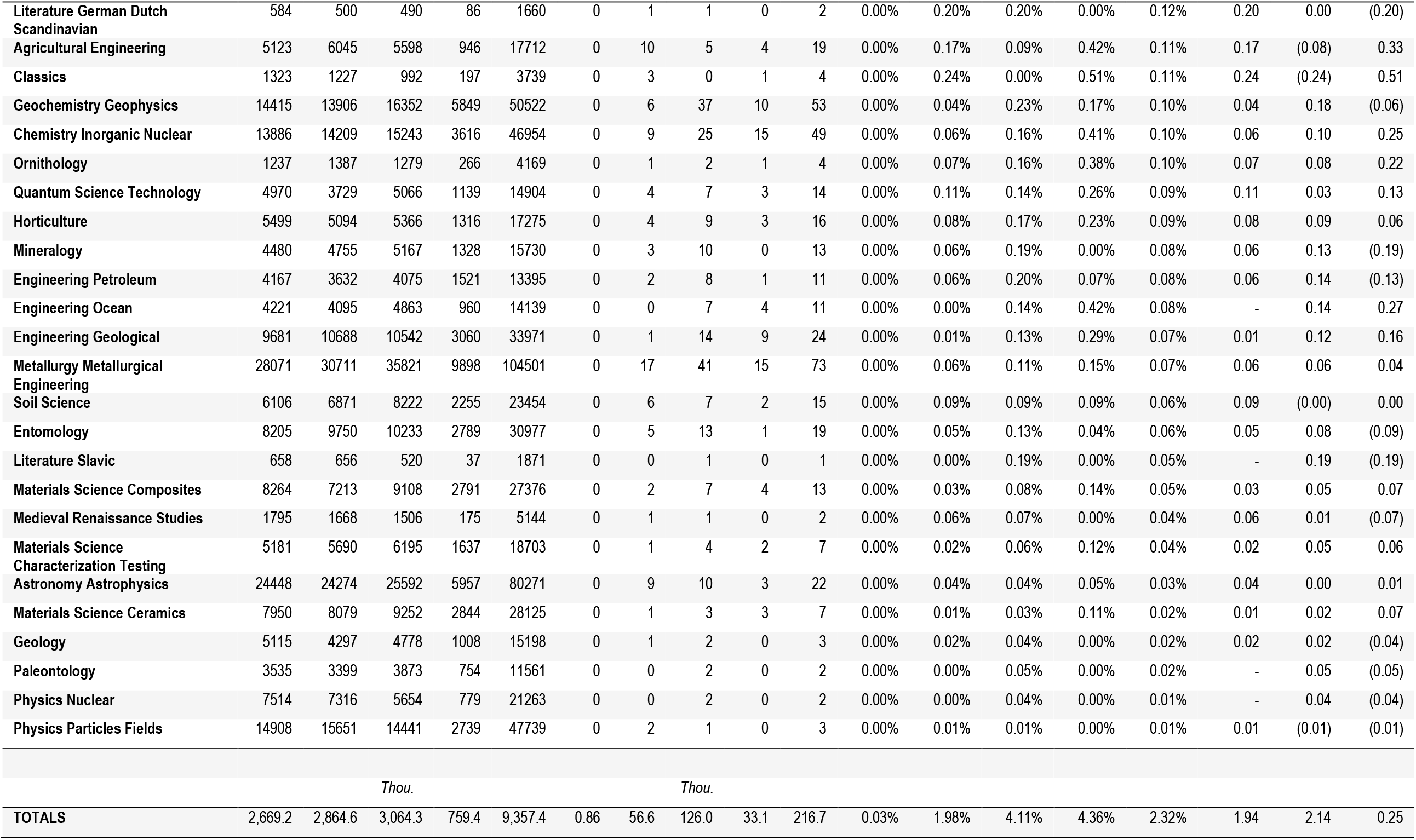
COVID 19 publications reported in the Web of Science (WoS), 2019-2022*. Source: See Note 1. * 2022 publications through to April 19, 2022

**Appendix 2. Table 3.**
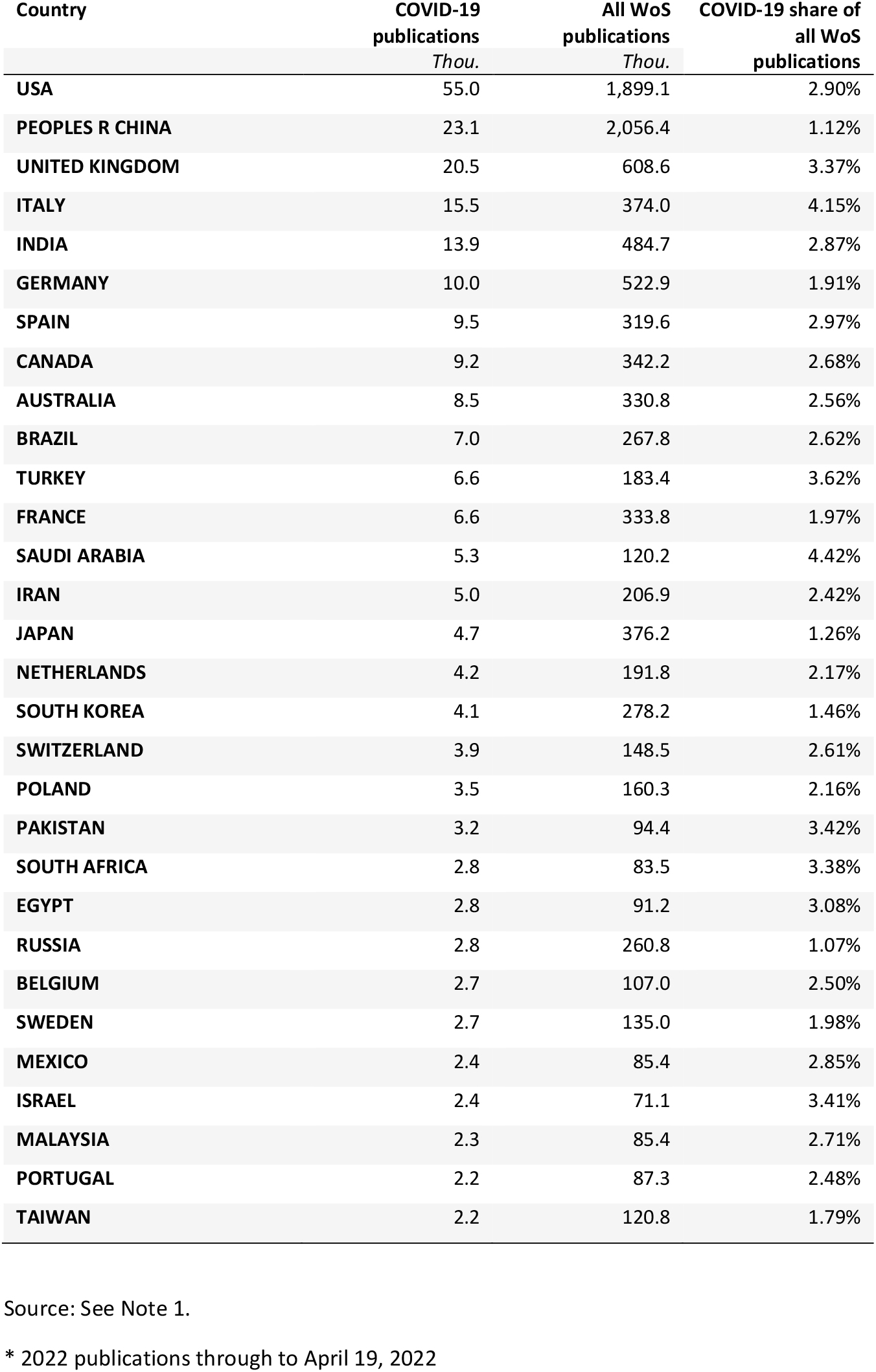
COVID-19 Publications by Top 30 Publishing Countries, Web of Science (WoS), 2019-2022*.

| Country | COVID-19<br>publications<br>*Thou.* | All WoS<br>publications<br>*Thou.* | COVID-19 share of<br>all WoS<br>publications |
| --- | --- | --- | --- |
| USA | 55.0 | 1,899.1 | 2.90% |
| PEOPLES R CHINA | 23.1 | 2,056.4 | 1.12% |
| UNITED KINGDOM | 20.5 | 608.6 | 3.37% |
| ITALY | 15.5 | 374.0 | 4.15% |
| INDIA | 13.9 | 484.7 | 2.87% |
| GERMANY | 10.0 | 522.9 | 1.91% |
| SPAIN | 9.5 | 319.6 | 2.97% |
| CANADA | 9.2 | 342.2 | 2.68% |
| AUSTRALIA | 8.5 | 330.8 | 2.56% |
| BRAZIL | 7.0 | 267.8 | 2.62% |
| TURKEY | 6.6 | 183.4 | 3.62% |
| FRANCE | 6.6 | 333.8 | 1.97% |
| SAUDI ARABIA | 5.3 | 120.2 | 4.42% |
| IRAN | 5.0 | 206.9 | 2.42% |
| JAPAN | 4.7 | 376.2 | 1.26% |
| NETHERLANDS | 4.2 | 191.8 | 2.17% |
| SOUTH KOREA | 4.1 | 278.2 | 1.46% |
| SWITZERLAND | 3.9 | 148.5 | 2.61% |
| POLAND | 3.5 | 160.3 | 2.16% |
| PAKISTAN | 3.2 | 94.4 | 3.42% |
| SOUTH AFRICA | 2.8 | 83.5 | 3.38% |
| EGYPT | 2.8 | 91.2 | 3.08% |
| RUSSIA | 2.8 | 260.8 | 1.07% |
| BELGIUM | 2.7 | 107.0 | 2.50% |
| SWEDEN | 2.7 | 135.0 | 1.98% |
| MEXICO | 2.4 | 85.4 | 2.85% |
| ISRAEL | 2.4 | 71.1 | 3.41% |
| MALAYSIA | 2.3 | 85.4 | 2.71% |
| PORTUGAL | 2.2 | 87.3 | 2.48% |
| TAIWAN | 2.2 | 120.8 | 1.79% |

Source: See Note 1.
\* 2022 publications through to April 19, 2022

**Appendix 2. Table 4.** COVID 19 publications (including preprints) reported in Dimensions, 2019-2022*.

| Dimensions research field | Field code | All publications<br>*Thou.* |  |  |  | COVID-19 publications<br>*Thou.* |  |  |  | COVID-19 share of all publications |  |  |  | Research pivot points |  |  |
| --- | --- | --- | --- | --- | --- | --- | --- | --- | --- | --- | --- | --- | --- | --- | --- | --- |
|  |  | 2019 | 2020 | 2021 | 2022* | 2019 | 2020 | 2021 | 2022* | 2019 | 2020 | 2021 | 2022* | 19-20 | 20-21 | 21-22* |
| Medical and Health Sciences | 11 | 1240.5 | 1463.4 | 1526.2 | 403.7 | 0.7 | 113.2 | 149.5 | 37.9 | 0.05% | 7.74% | 9.79% | 9.38% | 7.68 | 2.06 | -0.41 |
| Studies in Human Society | 16 | 162.8 | 192.1 | 206.5 | 48.9 | 0.0 | 10.2 | 17.9 | 4.4 | 0.01% | 5.28% | 8.69% | 8.97% | 5.27 | 3.40 | 0.28 |
| Biological Sciences | 6 | 358.4 | 406.9 | 439.3 | 127.0 | 0.4 | 8.5 | 11.6 | 3.2 | 0.10% | 2.09% | 2.63% | 2.51% | 1.99 | 0.54 | -0.12 |
| Psychology and Cognitive Sciences | 17 | 181.4 | 197.6 | 220.4 | 56.8 | 0.0 | 7.8 | 15.3 | 3.9 | 0.01% | 3.96% | 6.94% | 6.81% | 3.95 | 2.99 | -0.13 |
| Information and Computing Sciences | 8 | 373.8 | 408.0 | 450.4 | 112.1 | 0.0 | 6.7 | 14.1 | 3.5 | 0.01% | 1.65% | 3.12% | 3.11% | 1.65 | 1.47 | -0.02 |
| Economics | 14 | 92.2 | 104.5 | 111.1 | 28.8 | 0.0 | 5.8 | 9.3 | 2.3 | 0.02% | 5.54% | 8.36% | 7.83% | 5.53 | 2.82 | -0.54 |
| Education | 13 | 102.7 | 120.9 | 135.6 | 32.5 | 0.0 | 5.8 | 15.5 | 4.0 | 0.01% | 4.77% | 11.41% | 12.28% | 4.75 | 6.64 | 0.87 |
| Commerce, Management, Tourism and Services | 15 | 104.5 | 118.2 | 132.0 | 35.2 | 0.0 | 4.0 | 10.4 | 3.0 | 0.00% | 3.40% | 7.85% | 8.58% | 3.39 | 4.45 | 0.74 |
| Engineering | 9 | 660.6 | 700.6 | 762.8 | 251.5 | 0.0 | 2.9 | 6.0 | 1.8 | 0.00% | 0.42% | 0.79% | 0.70% | 0.42 | 0.37 | -0.08 |
| Mathematical Sciences | 1 | 215.0 | 231.3 | 247.0 | 70.5 | 0.0 | 2.8 | 4.0 | 1.1 | 0.00% | 1.20% | 1.64% | 1.54% | 1.19 | 0.44 | -0.10 |
| Chemical Sciences | 3 | 324.2 | 348.5 | 368.8 | 124.4 | 0.0 | 2.1 | 4.4 | 1.4 | 0.01% | 0.59% | 1.18% | 1.11% | 0.58 | 0.59 | -0.07 |
| Law and Legal Studies | 18 | 49.7 | 57.4 | 62.9 | 11.5 | 0.0 | 2.0 | 3.1 | 0.5 | 0.00% | 3.51% | 4.96% | 4.63% | 3.51 | 1.45 | -0.32 |
| History and Archaeology | 21 | 83.9 | 96.3 | 97.5 | 20.0 | 0.0 | 1.3 | 2.0 | 0.4 | 0.00% | 1.37% | 2.03% | 1.84% | 1.36 | 0.66 | -0.19 |
| Language, Communication and Culture | 20 | 100.7 | 113.9 | 118.8 | 25.5 | 0.0 | 1.3 | 2.8 | 0.7 | 0.00% | 1.14% | 2.33% | 2.89% | 1.13 | 1.19 | 0.56 |
| Philosophy and Religious Studies | 22 | 57.9 | 62.6 | 65.6 | 14.2 | 0.0 | 1.2 | 1.7 | 0.3 | 0.00% | 1.86% | 2.61% | 2.19% | 1.86 | 0.75 | -0.42 |
| Physical Sciences | 2 | 243.9 | 251.1 | 266.1 | 73.9 | 0.0 | 1.0 | 2.5 | 0.5 | 0.01% | 0.41% | 0.95% | 0.69% | 0.41 | 0.54 | -0.26 |
| Technology | 10 | 99.2 | 99.9 | 102.3 | 26.6 | 0.0 | 1.0 | 1.7 | 0.5 | 0.01% | 0.97% | 1.65% | 1.72% | 0.95 | 0.68 | 0.07 |
| Environmental Sciences | 5 | 79.8 | 91.6 | 110.9 | 33.6 | 0.0 | 0.9 | 2.1 | 0.6 | 0.01% | 1.00% | 1.91% | 1.70% | 0.99 | 0.90 | -0.21 |
| Agricultural and Veterinary Sciences | 7 | 72.5 | 82.7 | 90.7 | 27.6 | 0.1 | 0.9 | 1.2 | 0.2 | 0.17% | 1.04% | 1.34% | 0.87% | 0.87 | 0.30 | -0.47 |
| Built Environment and Design | 12 | 30.4 | 35.9 | 42.6 | 14.4 | 0.0 | 0.8 | 2.4 | 0.9 | 0.00% | 2.17% | 5.52% | 6.09% | 2.17 | 3.35 | 0.58 |
| Earth Sciences | 4 | 92.4 | 100.0 | 112.6 | 36.0 | 0.0 | 0.4 | 0.8 | 0.2 | 0.00% | 0.38% | 0.67% | 0.61% | 0.38 | 0.29 | -0.06 |
| Studies in Creative Arts and Writing | 19 | 20.0 | 21.1 | 22.5 | 4.7 | 0.0 | 0.3 | 0.8 | 0.2 | 0.02% | 1.48% | 3.47% | 3.49% | 1.46 | 1.99 | 0.02 |
| Totals |  | 5812.4 | 6535.8 | 6541.7 | 1681.4 | 1.3 | 207.7 | 309.4 | 75.7 | 0.02% | 3.18% | 4.73% | 4.50% | 3.16 | 1.55 | -0.23 |

Source: See Note 2. \* 2022 publications through to April 18, 2022

## Footnotes

1 WHO Director-General’s statement on IHR Emergency Committee on Novel Coronavirus (2019-nCoV). 30 January 2020. World Health Organization. https://www.who.int/dg/speeches/detail/who-director-general-s-statement-on-ihr-emergency-committee-on-novel-coronavirus-(2019-ncov).

2 WHO Director-General’s opening remarks at the media briefing on COVID-19 - 11 March 2020. World Health Organization. https://www.who.int/dg/speeches/detail/who-director-general-s-opening-remarks-at-the-media-briefing-on-covid-1911-march-2020.

3 Our World in Data, COVID-19 dataset, Oxford Martin Programme on Global Development, University of Oxford, and the Global Change Data Lab. https://github.com/owid/covid-19-data/tree/master/public/data (downloaded November 28, 2020).

4 Calculated from Our World in Data, COVID-19 dataset, ibid. Reference to countries also includes territories and dependencies separately reporting COVID-19 cases.

5 Hale, T., Boby, T., Angrist, N., Cameron-Blake, E., Hallas, L., Kira, B., Majumdar, S., Petherick, A., Phillips, T., Tatlow, H, and Webster, S. Variation in Government Responses to COVID19. Version 9.0. Blavatnik School of Government Working Paper. 24 November 2020. https://www.bsg.ox.ac.uk/sites/default/files/2020-11/BSG-WP-2020-032-v9.pdf (accessed November 28, 2020).

6 Our World in Data, OxCGRT COVID-19 Government Response dataset, https://github.com/owid/covid-19-data/tree/master/public/data/bsg (accessed November 28, 2020).

7 OECD, Draft summary of the STIO-GSF Virtual workshop on “Mobilising science in response to COVID-19,” 21 October 2020. Directorate for Science, Technology and Innovation, Committee for Scientific and Technological Policy, DSTI/STP/GSF/M(2020)2/ANN, Organisation for Economic Co-operation and Development, Paris. https://community.oecd.org/docs/DOC-184320 (accessed November 28, 2020).

8 Lem, P. Grant ‘repurposing’ towards Covid-19 widespread. Research Professional News, August 10, 2020. https://www.researchprofessionalnews.com/rr-news-europe-universities-2020-8-grant-repurposing-towards-covid-19-widespread/

9 Kaiser, J. NIH grapples with rush to claim billions in pandemic research funds. Science, June 3, 2020. http://doi.org/10.1126/science.abd1508.

10 OECD, COVID19 Research funding worldwide (to 21 September 2020). Organisation for Economic Co-operation and Development, Paris. https://community.oecd.org/docs/DOC-171875 (accessed November 28, 2020).

11 COVID-19 R&D Tracker. Policy Cures Research. https://www.policycuresresearch.org/covid-19-r-d-tracker (accessed November 28, 2020). This estimate is obtained from public funding announcements and press releases. Policy Cures Research notes that this is an incomplete estimate which does not include vaccine R&D and other R&D associated with the €7.4 billion ($8.8 billion) European Union Coronavirus Global Response and the more than $11.4 billion committed by the US Government to a range of agencies including BARDA (Biomedical Advanced Research and Development Authority). Chinese governmental COVID-19 R&D is not included, although the COVID-19 R&D Tracker does capture funding by some Chinese philanthropic and industrial organizations.

12 COVID-19 R&D Tracker, ibid.

13 Fry, C.V., Cai, X., Zhang, Y., and Wagner, C.S. Consolidation in a crisis: Patterns of international collaboration in early COVID-19 research. PLoS ONE, 2020, 15(7): e0236307. https://doi.org/10.1371/journal.pone.0236307.

14 Brainard, J. New tools to tame pandemic paper tsunami, Science, May 29, 2020, 368 (6494), 924-924.

15 Sharma, M., Scar, S., and Kelland, K. Coronavirus and the risks of ‘speed Science. 24 March 2020. World Economic Forum. https://www.weforum.org/agenda/2020/03/speed-science-coronavirus-covid19-research-academic (accessed October 23, 2020).

16 Torres-Salinas, D., Robinson-Garcia, N., Castillo-Valdivieso, P.A. Open Access and Altmetrics in the pandemic age: Forescast analysis on COVID-19 literature. bioRxiv, posted April 26, 2020. https://doi.org/10.1101/2020.04.23.057307.

17 Council on Governmental Relations. Research Impact under COVID-19. August 25, 2020. Washington, DC. https://www.cogr.edu/sites/default/files/Research_COVID_August2020_COGR_FINAL.pdf (accessed September 3, 2020).

18 Vitae. Covid-19 impact on researchers. Study commissioned by the Department for Business, Energy and Industrial Strategy, UK. October 12, 2020. Results reported at https://www.vitae.ac.uk/news/vitae-news-2020/impact-of-lockdown-on-researchers-in-uk (accessed November 29, 2020).

19 Larkins, F., et al., Impact of the pandemic on Australia’s research workforce. May 8, 2020. Research Brief. Australian Government, Chief Scientist. https://www.science.org.au/sites/default/files/rrif-covid19-research-workforce.pdf (accessed November 29, 2020).

20 European Commission, Impact of Covid-19 on International Students in EE and OECD Member States. Inform #2. https://ec.europa.eu/home-affairs/sites/homeaffairs/files/00_eu_inform2_students_final_en.pdf (accessed November 29, 2020).

21 Andersen, J.P., Nielsen, M.W., Simone, N.L., Lewiss, R.E., and Jagsi, R. Meta-Research: COVID-19 medical papers have fewer women first authors than expected. June 15, 2020. eLife 2020;9:e58807. https://doi.org/10.7554/eLife.58807.

22 Myers, K.R., Tham, W.Y., Yin, Y., Cohodes, N., Thursby, J.G., Thursby, M.C., Schiffer, P., Walsh, J.T., Lakhani, K.R., and Wang, D. Unequal effects of the COVID-19 pandemic on scientists. Nature Human Behaviour, 4, 880–883 (2020). https://doi.org/10.1038/s41562-020-0921-y.

23 Blanding, M. Research in the Time of COVID. Technology Review. October 20, 2020. https://www.technologyreview.com/2020/10/20/1009361/research-in-the-time-of-covid/ (accessed October 24, 2020).

24 Institute for Cancer Research. Pandemic to delay cancer advances by nearly 18 months, researchers fear. November 30, 2020, London, https://www.icr.ac.uk/news-archive/pandemic-to-delay-cancer-advances-by-nearly-18-months-researchers-fear (accessed November 30, 2020).

25 Ball, P. Why lockdown silence was golden for science. The Guardian, 20 June 2020. https://www.theguardian.com/science/2020/jun/20/why-lockdown-silence-was-golden-for-science (accessed June 21, 2020).

26 Stokstad, E. Pandemic lockdown stirs up ecological research, Science, 28 August 2020, 369 (6506), 893.

27 Grim, D. As labs move to reopen, safety worries abound. Science, 15 May 2020. 368 (6492), 690-691.

28 JASON. Managing the Risk from COVID-19 During a Return to On-Site University Research. JSR-20-NS1. August 25, 2020. MITRE Corporation, VA.

29 Working safely during coronavirus (COVID-19). Labs and research facilities. Department for Business, Energy & Industrial Strategy and Department for Digital, Culture, Media & Sport, UK. Updated 27 November 2020. https://www.gov.uk/guidance/working-safely-during-coronavirus-covid-19/labs-and-research-facilities (accessed November 29, 2020).

30 Institute for Cancer Research, op. cit. (footnote 24).

31 As of November 29, 2020, PubMed reports a total of 31,813,673 publication records (https://pubmed.ncbi.nlm.nih.gov/?term=%221800%3A2100%5Bdp%5D%22). Of these, 8,371,581 are free full text (https://pubmed.ncbi.nlm.nih.gov/?term=%221800%3A2100%5Bdp%5D%22&filter=simsearch2.ffrft) from NLMs PubMed Central (full-text database of articles) and other sources.

32 https://clarivate.com/webofsciencegroup/solutions/web-of-science-core-collection/. A search of the Web of Science (November 29, 2020) returns 78,153,527 publication records (Indexes=SCI-EXPANDED, SSCI, A&HCI, CPCI-S, CPCI-SSH, BKCI-S, BKCI-SSH, ESCI, CCR-EXPANDED)

33 Publications in languages other than English comprised 14.9% and 7.5% respectively of the total number of PubMed and WoS reported in footnotes 31 and 32.

34 http://www.techminingforglobalgood.org/open-covid-19-research-for-analysis/.

35 http://www.techminingforglobalgood.org/open-covid-19-research-for-analysis/ (accessed October 24, 2020).

36 See: https://www.thevantagepoint.com/.

37 In 2000, there were 141 WoS and 151 PubMed publication recorded on coronavirus-related topics.

38 Organisation for Economic Cooperation and Development.

39 Comparison of COVID-19 WoS study dataset with aggregate WoS publications analyzed by countries (same indexes and document types) for January 1, 2020 -November 30, 2020 (N=2,199,639).

40 In the top 100 COVID-19 publishing organizations by author affiliations in the PubMed dataset, 54 are indicated as universities, 25 as medical schools, 18 are hospitals or clinics, and 3 are national or public research organizations. However, organizations and affiliations can be intertwined. Hospitals and clinics are often associated with universities. In some cases, researchers will publish with a university or medical school or hospital affiliation (or a combination of two or more affiliations).

41 Wuhan is the city where the first large scale outbreak of COVID-19 was identified. See: Novel Coronavirus – China, 12 January 2020, World Health Organization, https://www.who.int/csr/don/12-january-2020-novel-coronavirus-china/en/; C. Huang et al., Clinical features of patients infected with 2019 novel coronavirus in Wuhan, China, The Lancet, January 24, 2020, https://doi.org/10.1016/S0140-6736(20)30183-5.

42 While there are broad similarities in outputs of COVID-19 PubMed and WoS papers by organizational affiliations, detailed differences are also observable. Underlying reasons for these differences may include variations in organizational structure (e.g., university relationships with medical schools and affiliated hospitals) and disciplinary composition within each institution, topical foci, and distribution of publication outlets including by journal, peer reviewed and preprint placements.

43 Web of Science Core Collection: Web of Science Categories. Clarivate. https://support.clarivate.com/ScientificandAcademicResearch/s/article/Web-of-Science-Core-Collection-Web-of-Science-Categories?language=en_US (accessed December 3, 2020).

44 See: Milojević, S. Practical method to reclassify Web of Science articles into unique subject categories and broad disciplines. Quantitative Science Studies 2020 1:1, 183-206. https://doi.org/10.1162/qss_a_00014.

45 Wang, Q., and Waltman, L. Large-scale analysis of the accuracy of the journal classification systems of Web of Science and Scopus. Journal of Infometrics, 2020, 10, 347-364. https://doi.org/10.1016/j.joi.2016.02.003.

46 The 2017-2019 period was chosen to provide a recent comparison and to gather a reasonably sized dataset (N=2,303) across WoS SCs.

47 Percentage totals by SCs add up to more than 100% reflecting articles in journals that are assigned two or more SCs.

48 Björk, B-C., and Solonon, D. The publishing delay in scholarly peer-reviewed journals. Journal of Infometrics, 2020, 7, 914-923, http://dx.doi.org/10.1016/j.joi.2013.09.001. Note: with the growth of preprints and early access, papers are increasingly publicly available and citable prior to official publication, see, for example: Haustein, S., Bowman, R., and Costas, R. When is an article actually published? An analysis of online availability, publication, and indexation dates. arXiv: 1505.00796, 2015. https://arxiv.org/abs/1505.00796.

49 Journal peer review processes in 2020 have also been affected by the pandemic. Journals dealing with COVID-19 topics have seen huge increases in submissions, with peer reviewers under pressure to review quickly, while in other journal fields, editorial and review delays are reported. Elsevier, Review delays during coronavirus crisis,

50 As reported in Figure 2.

51 The aggregated WoS publication numbers reported in this analysis are slightly lower than reported in Figure 2. The reason is that the subject category analysis in this section was undertaken with data as of October 24, 2020 (when WoS reported 1.95 million records), while Figure 2 is based on data from November 30, 2020 (when WoS reported 2.105 million records) 2020 (Indexes=SCI-EXPANDED, SSCI, A&HCI, CPCI-S, CPCI-SSH, ESCI) refined by document types (article, early access, review, or proceedings paper).

52 The 2020 COVID-19 share of all WoS publications is lower than reported for PubMed (see earlier discussion and Figure 2). PubMed focus on medical, bioscience, and public health domains and includes preprints, while WoS primarily covers journals across all fields of science, social science, and the arts and humanities.

53 These fields vary in size, with total SC publishing (in 2020) ranging from many thousands to a few hundred. While “Folklore” is the smallest field in the top quintile (*RP*=3.8, *cPC19*=0.0%; *cPC20*=3.8%), researchers in this field have published in 2020 on such subjects as myths and urban legends about coronavirus, and how the pandemic has affected museums and traditional folk crafts.

54 SCI=Science Citation Index; SSCI = Social Science Citation Index; A&HCI = Arts and Humanities Citation Index.

